# A *Toxoplasma gondii* O-glycosyltransferase that modulates bradyzoite cyst wall rigidity is structurally and functionally distinct from host homologues

**DOI:** 10.1101/2023.10.26.564164

**Authors:** Pranav Kumar, Tadakimi Tomita, Thomas A. Gerken, Collin J. Ballard, Yong Sok Lee, Louis M. Weiss, Nadine L. Samara

## Abstract

Infection with the Apicomplexan protozoan *Toxoplasma gondii* can be life-threatening in immunocompromised hosts. Transmission frequently occurs through the oral ingestion of *T. gondii* bradyzoite cysts, which transition to tachyzoites, disseminate, and then form cysts containing bradyzoites in the central nervous system, resulting in latent infection. There are currently no effective treatments to cure latent infection. Bradyzoites are encapsulated by a cyst wall that is critical for immune evasion, survival, and transmission. Cyst wall rigidity is influenced by the O-glycosylation of the mucin domain of the cyst wall protein CST1 by mucin-type O-glycosyltransferases (GalNAc-Ts). Here, we report the first structures of a protozoan GalNAc-T, *T.gondii*-GalNAc-T3 in the apo state and in complex with glycopeptide substrates. The structures reveal features that are strictly conserved in Apicomplexan homologues of *T.gondii*-GalNAc-T3, including a unique 2^nd^ metal binding site that is coupled to substrate binding and enzymatic activity *in vitro* and cyst wall O-glycosylation in *T. gondii*. Additional structural features illustrate the divergence of GalNAc-Ts from parasite to host and highlight multiple druggable sites in *T.gondii*-GalNAc-T3 and its homologues in Apicomplexa that are responsible for a wide range of parasitic diseases.

## Introduction

*Toxoplasma gondii* is an obligate intracellular protozoan pathogen belonging to the phylum Apicomplexa that infects a wide range of warm-blooded animals, including humans^1^. It is transmitted through the oral ingestion of bradyzoites found in tissue cysts in muscle or brain (i.e. undercooked meat) or by the ingestion of sporozoites found in oocysts (found in cat fecal matter) contaminating food or water^2^. *T. gondii* tissue cysts contain bradyzoites, reside in the central nervous system or muscle tissue, and are associated with latent infection. These tissue cysts are the source of recrudescent infection in immunocompromised hosts. During recrudescent infection, bradyzoites convert to tachyzoites and disseminate resulting in infections that can be life threatening in immunocompromised hosts such as HIV/AIDS patients^3^. While there are drugs that target tachyzoites, *T. gondii* tissue cysts have thus far been resistant to treatment^4^. Thus, finding an effective medication that targets bradyzoites in tissue cysts is critical for preventing infection and disease.

The *T. gondii* tissue cyst is surrounded by a wall that is critical for the survival and transmission of *T. gondii*. The cyst wall contains a subset of proteins that undergo mucin-type O-glycosylation, a post-translational modification (PTM) that is conserved across eukaryotes and occurs on proteins that pass through the secretory pathway^5^. *T. gondii* O-glycosylated cyst wall proteins include CST1 (TGME49_264660)^6^, SRS13 (TGME49_222370)^7^, proteophosphoglycan PPG1 (TGME49_297520)^6,8^, and GRA2 (TGME49_227620)^9^. CST1 is an abundant protein that is critical for maintaining cell wall integrity and promoting bradyzoite persistence. It contains thirteen SRS repeats, a mucin domain with 20 threonine-rich repeats, and a C-terminal cysteine-rich domain (Fig. 1a). O-glycosylation of CST1 occurs within the mucin domain and is critical for its function in the formation of an organized and structurally stable cyst wall^6,10^.

**Fig. 1:**
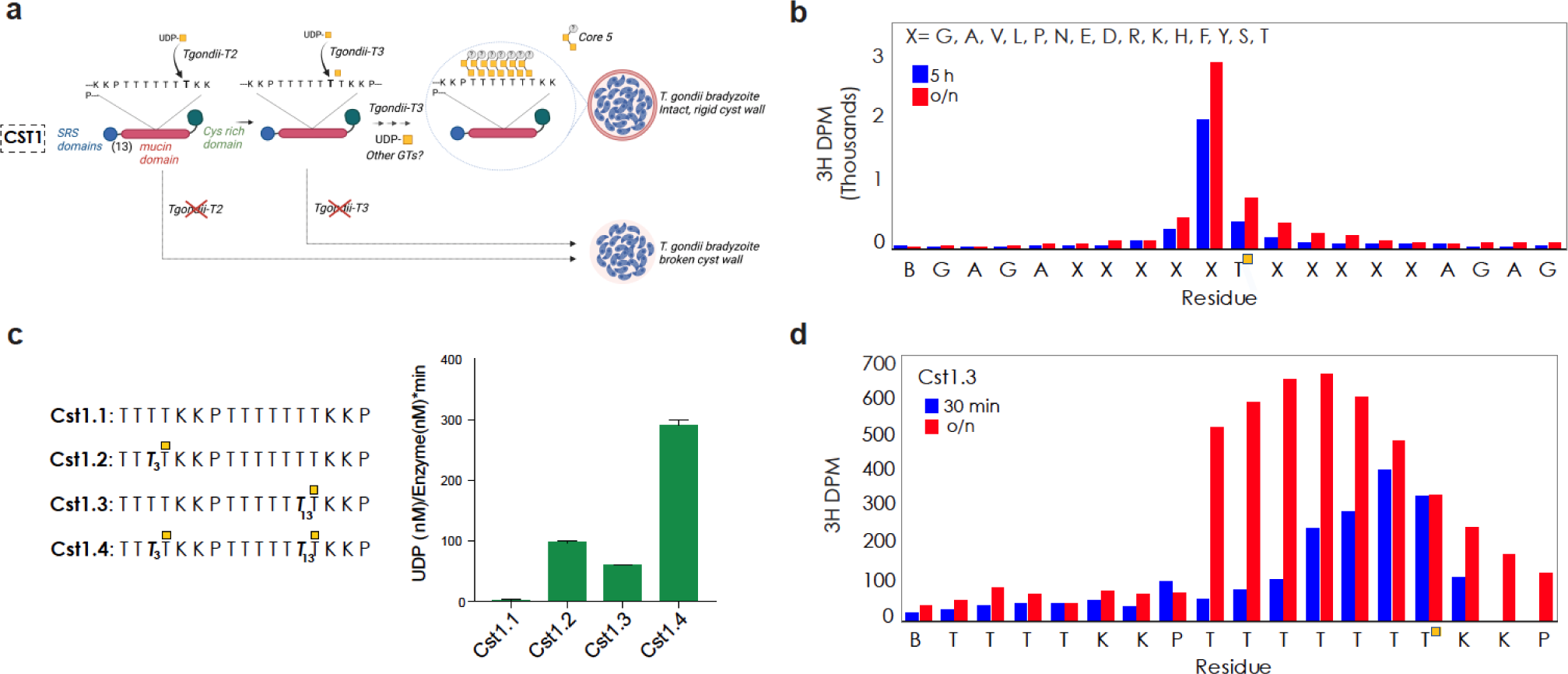
Mucin-type O-glycosylation by *T.gondii*-GalNAc-T3. **a,** O-glycosylation of the mucin domain of cyst wall protein CST1 by *T.gondii*-GalNAc-T2 and *T.gondii*-GalNAc-T3 occurs in a hierarchal manner. *T.gondii*-GalNAc-T2 initiates the process by O-glycosylating a unglycosylated acceptor Thr, and *T.gondii*-GalNAc-T3 recognizes a Thr-O-GalNAc at the +1 position to glycosylate adjacent sites in the N-terminal direction. O-glycosylation of CST1 influences cyst wall rigidity. Knocking out each transferase results in a fragile cyst wall. Figure created with BioRender.com. **b,** Edman degradation assay using the GPIIC random library shows that *T.gondii*-GalNAc-T3 specificity is dictated by GalNAc at the +1 position in a sequence independent manner. **c,** *T.gondii*-GalNAc-T3 O-glycosylation of Cst1 glyco-peptides increases with additional Thr-O-GalNAcs. **d,** Edman degradation assay shows that O-glycosylation by *T.gondii*-GalNAc-T3 is processive after 30 min, with decreasing efficiency after each glycosylation. *T.gondii*-GalNAc-T3 densely glycosylates the Cst1.3 mono-glycopeptide after an overnight reaction.

Mucin-type O-glycosylation is initiated by a large family of GalNAc-Ts, which catalyze the transfer of N-acetylgalactosamine (GalNAc) from UDP-GalNAc to Thr/Ser in a Mn^2+^ dependent manner to form GalNAcα-O-Thr/Ser^11^. GalNAc-Ts are Golgi membrane-anchored enzymes that contain a catalytic domain (CAZy family GT27) that adopts a GT-A fold, and are the only known glycosyltransferases with a C-terminal Ricin B-type lectin domain^12^. *T. gondii* has 5 GalNAc-Ts^13–15^: *T.gondii*-GalNAc-T1, *T.gondii*-GalNAc-T2, and *T.gondii*-GalNAc-T3 are expressed in the bradyzoite and tachyzoite stages^10^, and *T.gondii*-GalNAc-T4 and *T.gondii*-GalNAc-T5 are expressed in the oocyst stage^10,16^. The role of mucin-type O-glycosylation in tachyzoites and oocysts is currently not known. In bradyzoites, *T.gondii*-GalNAc-T2 initiates the dense O-glycosylation of the Thr-rich mucin domain of CST1 by modifying unglycosylated regions. The strict glycopeptide-preferring enzyme *T.gondii*-GalNAc-T3 can then add GalNAc to an acceptor Thr one position N-terminal to a Thr-O-GalNAc and densely O-glycosylate stretches of Thr. The initial GalNAcs are then extended to form a core 5 O-glycan with an unknown capping sugar (Fig. 1a)^10^. Knocking out *T.gondii*-GalNAc-T2 or *T.gondii*-GalNAc-T3 results in cyst wall breakage, indicating that hierarchical and sequential glycosylation is critical for the ability of CST1 to generate a rigid cyst wall (Fig. 1a)^10^.

Reduced sequence homology between *T.gondii*-GalNAc-T3 and metazoan homologues hints that an evolutionary divergence may have occurred in substrate recognition and enzyme function, supporting the possibility that *T.gondii*-GalNAc-T3 could be specifically inhibited as a target for therapy to treat disease. To gain insight into the structural and functional differences between this enzyme and its human homologues, we solved X-ray crystal structures of *T.gondii*-GalNAc-T3 alone or in complex with glycopeptide substrates. The structures reveal a unique GalNAc binding pocket that dictates the substrate preference of *T.gondii*-GalNAc-T3 and is coupled to a 2^nd^ metal binding site that is strictly conserved among its Apicomplexan homologues. We also identify an active site Glu that is oriented for the deprotonation of the acceptor Thr. In addition, *T.gondii*-GalNAc-T3 has an extended C-terminal tail that is critical for its function, and a substrate binding loop that is absent in metazoan isoenzymes. Biochemical, computational, and *in vivo T. gondii* cyst wall glycosylation assays show that these unique structural features are critical for the enzymatic function of *T.gondii*-GalNAc-T3 and set a framework for potential sites of inhibition.

## Results

### *T. gondii*-GalNAc-T3 glycosylates N-terminal to a prior Thr-O-GalNAc in a sequence independent manner

The luminal region (catalytic and lectin domains, aa 74–635) of *T.gondii*-GalNAc-T3 (AY160970.1) was produced by secreted expression in *Pichia pastoris.* We first expanded on earlier studies showing that *T.gondii*-GalNAc-T3 glycosylates one residue N-terminal to a prior site (+1 position) using various substrates^14^. With the Muc5AC mucin (glyco)peptide series, *T.gondii*-GalNAc-T3 does not efficiently glycosylate the nascent Muc5AC peptide, but readily glycosylates the mono-glycopeptides Muc5AC-3 and Muc5AC-13 and the di-glycopeptide Muc5AC-3,13 (Fig. S1a). Using a random glycopeptide substrate library (GPIIC)^17^, we observe that *T.gondii*-GalNAc-T3 activity is enhanced by Thr-O-GalNAc at the +1 position of an acceptor site independent of the surrounding amino acid sequence (−5 to +5 aa, Fig. 1b).

We verified the substrate preference of *T.gondii*-GalNAc-T3 using peptides and glycopeptides from *T. gondii* cyst wall proteins CST1 and SRS13. For CST1, *T.gondii*-GalNAc-T3 does not efficiently glycosylate the unglycosylated peptide Cst1.1, is similarly active towards the mono-glycopeptides Cst1.2 and Cst1.3 and has the highest activity against the di-glycopeptide Cst1.4, as expected (Fig. 1c). *T.gondii*-GalNAc-T3 does not readily modify the unglycosylated Srs13.1 peptide, as predicted (Fig. S1b). Surprisingly, there is diminished activity for the C-terminal mono-glycopeptide Srs13.3, while activity for Srs13.4 di-glycopeptide is ∼1.5-fold lower than the Srs13.2 mono-glycopeptide despite the availability of 2 potential acceptor sites, suggesting that either the C-terminal acceptor site or proximal amino acids are diminishing enzymatic activity. Differences in activity among the various peptide series suggest that while proximal peptide sequence does not affect the preference for Thr-O-GalNAc at +1, it influences the efficiency at which *T.gondii*-GalNAc-T3 can modify an acceptor.

Earlier studies indicated the mucin domain of CST1 was fully glycosylated ^10^. We thus assessed the ability of *T.gondii*-GalNAc-T3 to fully glycosylate glycopeptide substrates. For the CST1 glycopeptides, *T.gondii*-GalNAc-T3 sequentially glycosylates sites N-terminal to the Thr-O-GalNAc with decreasing efficiency upon each GalNAc addition in 30-or 90-min reactions (Fig. 1d, Fig. S1c) and can eventually fully glycosylate all 6 N-terminal Thr on Cst1.3 in an overnight reaction (Fig. 1d and Fig. S1g). We observe similar sequential and gradually decreasing glycosylation efficiency for the Muc5Ac glycopeptides and Srs13.2 (Fig. S1d,e). The decreasing efficiency upon each GalNAc addition suggests that *T.gondii*-GalNAc-T3 O-glycosylates its substrates using a distributive mechanism, where the substrate is released after a single GalNAc is added and forms a unique enzyme substrate complex for each addition, with decreasing affinity for the substrate with each additional GalNAc^18^. Interestingly, *T.gondii*-GalNAc-T3 efficiently and sequentially glycosylates Thr then Ser on Muc5Ac-13, and then there is a striking decrease in the efficiency of glycosylation at the sites N-terminal to Ser, suggesting that *T.gondii*-GalNAc-T3 specifically prefers Thr-O-GalNAc over Ser-O-GalNAc at the +1 position (Fig. S1d). Indeed, changing Thr-O-GalNAc to Ser-O-GalNAc on a glycopeptide results in a 5-fold increase in the K_M_, demonstrating diminished binding to Ser-O-GalNAc (Fig. S1f). Nevertheless, the overnight incubation of Muc5AC-13 with *T.gondii*-GalNAc-T3 will fully glycosylate all four acceptors in the TTSTT* sequence as shown by the Edman sequencing chromatograms (Fig. S1g).

### X-ray crystal structures of *T.gondii*-GalNAc-T3 reveal a non-conserved 2^nd^ metal binding motif that modulates substrate binding and regulates catalysis

To understand the basis of the substrate specificity and function of *T.gondii*-GalNAc-T3, we solved the X-ray co-crystal structures of *T.gondii*-GalNAc-T3 in complex with Mn^2+^, a non-hydrolysable form of UDP-GalNAc (UDP-2-(acetylamino)-4-F-D-galactosamine disodium salt, UDP-GalNAc-F) and each of the following glycopeptides: Cst1.4, Muc5AC-3,13, Muc5AC-3, Muc5AC-13, and Srs13.2 from 2.2-2.9 Å resolution (Fig. 2a, Fig. S2a-d, Table S1). The structures reveal a similar overall fold to metazoan homologues (Fig. S3a) where the active site adopts the conserved GT-A fold configuration with a DXH motif (Asp276, Ser277, His278) and Asp276, His278, His414, UDP, and 1 water molecule coordinate Mn^2+^_A_ with octahedral geometry (Fig. 2b, Fig. S3b)^19–26^. We modeled UDP into the active site but did not include GalNAc-F due to weak electron density, indicating that hydrolysis of UDP-GalNAc-F may have occurred during crystallization (Fig. S3b).

**Fig. 2:**
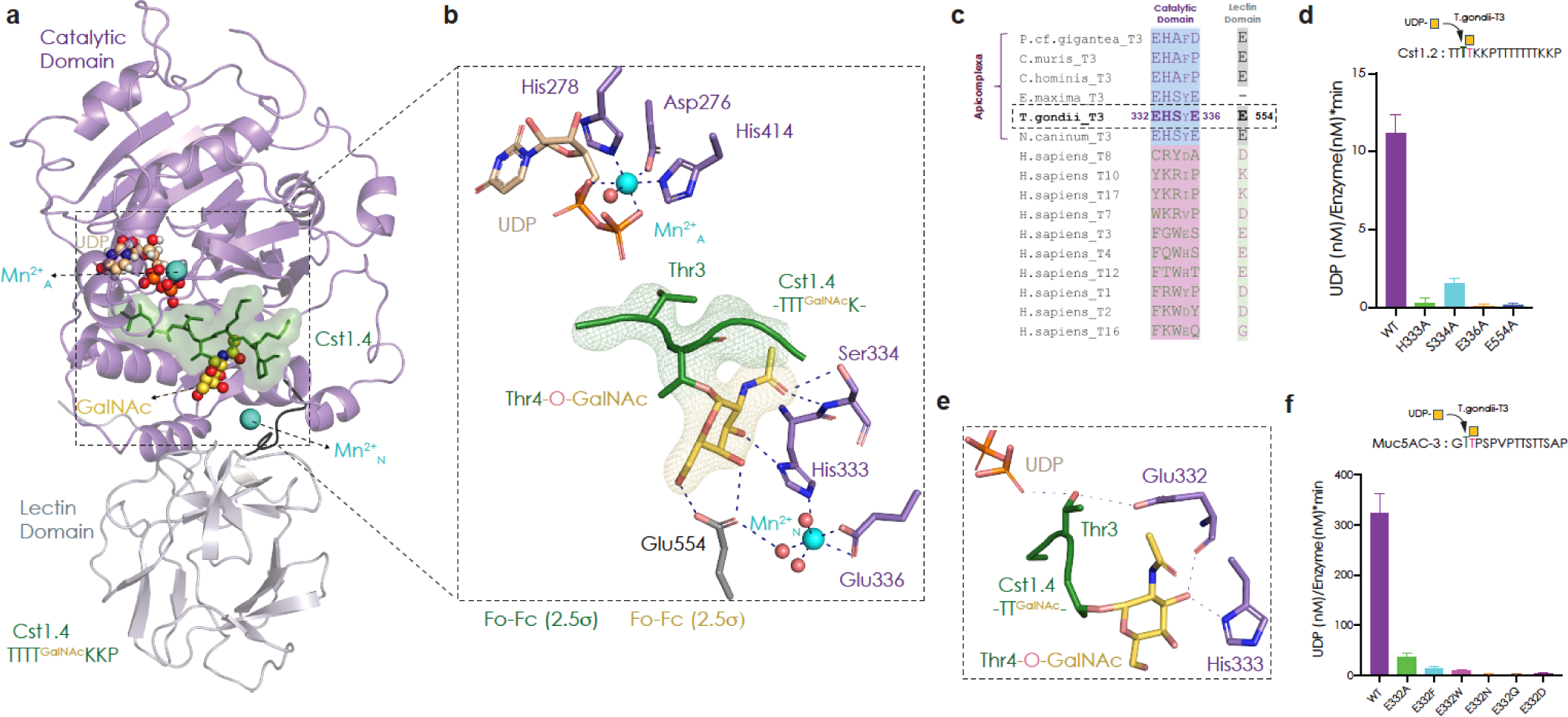
Co-crystal X-ray structure of *T.gondii*-GalNAc-T3, UDP, Mn^2+^, and a CST1 di-glycopeptide. **a,** *T.gondii*-GalNAc-T3, with the catalytic domain in lavender and the C-terminal lectin domain in grey. The Cst1.4 peptide is green, GalNAc is yellow. Mn^2+^ is cyan, and UDP is wheat. **b,** The active site adopts the conserved GT-A fold DXH motif. The acceptor Thr is correctly positioned for GalNAc transfer. GalNAc on the +1 Thr-O-GalNAc is bound to a pocket that is coupled to a 2^nd^ metal binding site (Mn^2+^_N_). Cst1.4 Fo-Fc omit map is shown in green and the GalNAc omit map is yellow, both contoured at 2.5σ. **c,** Sequence alignment comparing the GalNAc binding pocket residues of Apicomplexan homologues of *T.gondii*-GalNAc-T3 to a subset of representative human homologues showing the site is not conserved in higher eukaryotes. **d,** Activity assay comparing *T.gondii*-GalNAc-T3^WT^ to variants with disruptions in the 2^nd^ metal binding pocket showing that this site is important for enzymatic function, using Cst1.2 as a substrate. **e,** The unique residue Glu332 is positioned for acceptor Thr deprotonation. **f,** Activity assay showing that Glu332 influences enzymatic activity using Muc5A-3 as a substrate.

The N-terminal portions of the di-glycopeptides Cst1.4 (TTTT^GalNAc^KKP, Fig. 2a) and Muc5AC-3,13 (GTT^GalNAc^PSP, Fig. S2a) are ordered in the structure, but electron density for the remaining residues is weak. The peptides are similarly bound to the active site substrate groove with the acceptor Thr positioned for catalysis (Fig. 2a, Fig. S4a)^20^. For the mono-glycopeptides, we observe density for the substrate GalNAc in the Muc5AC-3 structure (Fig. S2b), but disconnected and weak density for the peptide residues. We could not model substrate density in the structure containing Muc5AC-13 due to unresolvable electron density, and we only observe weak density for GalNAc in the Srs13.2 structure (Fig. S2c and d). It is notable that the di-glycopeptides were the only substrates with strong electron density for the N-terminal portion of the peptide chain, but it is not clear what is stabilizing the peptide conformations in these structures since the glycopeptide C-termini were not resolvable.

Thr-O-GalNAc at the +1 position on the glycopeptides is bound to a pocket in *T.gondii*-GalNAc-T3 consisting of His333 (catalytic domain) and Glu554 (lectin domain) that interact with GalNAc via sidechain H-bonds, and Ser334 (catalytic domain), which interacts with GalNAc through a mainchain H-bond (Fig. 2b, S2a, b, S4a). A striking and unexpected feature of the GalNAc binding pocket is the presence of a 2^nd^ metal (Mn^2+^_N_) that has not previously been observed in metazoan GalNAc-Ts (Fig. S3a). Mn^2+^_N_ is coupled to the GalNAc binding pocket through coordination to His333 and indirect coordination to Glu554 via water. Glu336 and 2 additional waters complete the coordination sphere of Mn^2+^_N,_ which adopts an irregular octahedral geometry (Fig. 2b). The GalNAc and Mn^2+^_N_ binding residues are conserved in most Apicomplexan homologues of *T.gondii*-GalNAc-T3, and divergent from the corresponding region in human GalNAc-Ts, including GalNAc-T10, T7, and T17, which also strictly recognize GalNAc at the +1 position and modify the adjacent acceptor site (Fig. 2c)^17,27^. Disruption of the GalNAc pocket and Mn^2+^_N_ binding residues diminishes activity compared to *T.gondii*-GalNAc-T3^WT^, supporting a critical role for Mn^2+^_N_ in enzymatic function (Fig. 2d).

Interestingly, the GalNAc binding residue Glu332 within the metal binding motif is proximal to the acceptor Thr (Fig. 2e). In human isoenzymes, the residue at that position is frequently Arg, Trp, or Tyr (Fig. 2c). Changes in Glu to Ala or any of these residues abrogates activity, suggesting a unique key role for Glu332 in catalysis, possibly as a general base (Fig. 2f). Additionally, polar side chains, and Asp result in diminished activity, suggesting a strict preference for Glu332, which could help position the acceptor Thr for catalysis and/or increase the negative charge by deprotonating the Thr hydroxyl group. To further assess the role of Glu332 in enzymatic function, we performed enzyme kinetics on *T.gondii*-GalNAc-T3^WT^ and *T.gondii*-GalNAc-T3^E332A^ using Muc5AC-3 as a substrate. An ∼18-fold decrease in V_max_ and 5-fold decrease in K_M_ for *T.gondii*-GalNAc-T3^E332A^ compared to *T.gondii*-GalNAc-T3^WT^ supports a role for Glu332 in influencing reaction chemistry, by possibly aligning and/or deprotonating the acceptor Thr (Fig. S4b and c, Table S2).

### The 2^nd^ metal binding site regulates enzymatic function by influencing GalNAc binding and catalysis in a pH dependent manner

To examine the role of Mn^2+^_N_ in catalysis, we first verified its presence by solving the X-ray crystal structure of apo *T.gondii*-GalNAc-T3 in the absence of Mn^2+^ to 2.9 Å resolution and did not observe density for Mn^2+^_N_ (Fig. S5a, left panel, Table S3). We then solved the structure to 2.5 Å resolution after soaking the apo crystals in Mn^2+^ and detected the appearance of electron density and an anomalous signal for Mn^2+^_N_ (Fig. S5a, right panel, Table S3). We initially hypothesized that Mn^2+^_N_ was aligning the sidechains for GalNAc binding. However, a comparison of the substrate bound structure and apo structure shows that the Mn^2+^_N_ binding residues adopt similar conformations in both structures, indicating that Mn^2+^_N_ uses an alternate mechanism to influence activity (Fig. 3a).

**Fig. 3:**
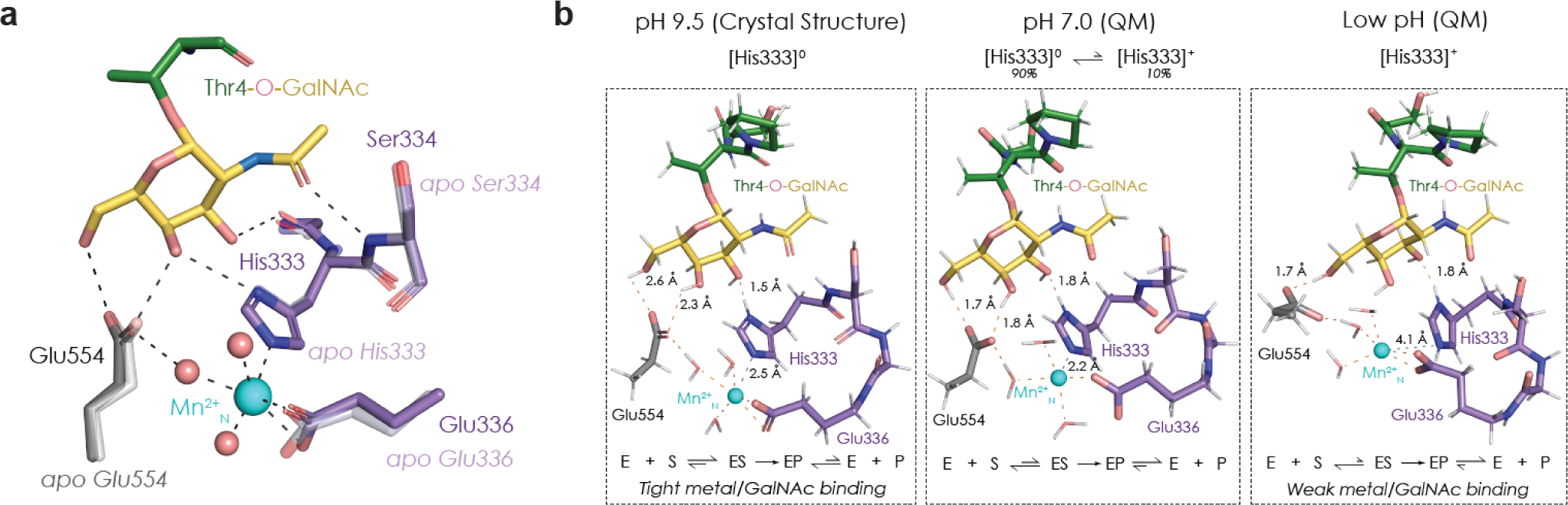
Effects of the second metal binding site on catalysis. **a,** There are no differences in the sidechain conformations of the metal binding residues in the apo structure (no Mn^2+^) and the co-crystal complex. **b,** The second metal influences catalysis by regulating Michaelis complex formation (ES) in a pH sensitive manner. **Left panel**, conformation of the site in the crystal structure at pH 9.5. Deprotonated sidechains bind tightly to Mn^2+^ and GalNAc, forming a tight Michaelis complex and inhibiting product release. **Middle panel**, quantum chemistry calculations at pH 7.0, where His deprotonation promotes ES complex formation, and His protonation promotes product release. **Right panel**, quantum chemistry calculations where the sidechains are fully protonated. Here, His333 is no longer binding Mn^2+^, and Glu554 interactions with GalNAc are reduced, disrupting ES complex formation and catalysis.

We then reasoned that the positive charge on Mn^2+^_N_ influences GalNAc binding and reaction chemistry. To assess the effect of electrostatics on activity, we began by measuring enzymatic activity at various pHs. Optimal activity occurs at around pH 7.3-7.8, and gradually decreases as pH increases (Fig. S5b). However, the crystals grow at pH 9.5, where activity is low in solution. Indeed, kinetics show that at pH 9.0, the V_max_ is 500-fold less than at pH 7.3 and the K_M_ is 4-fold less, suggesting tighter substrate binding but lower turnover (Fig. S5c). At pH 9-9.5, the acidic side chains are fully deprotonated and His333 is neutral and can coordinate Mn^2+^. Thus, the state observed in the crystal at pH 9.5 represents a possible stable E:S complex where substrate is tightly bound, and catalytic activity and turnover are reduced since GalNAc-Ts function optimally at around neutral pH (Fig. 3b, left panel)^28^. We could not obtain diffracting crystals at neutral pH, so we soaked crystals grown at pH 9.5 into a pH 7.3 buffer before freezing and did not observe bound Muc5AC-3,13 diglycopeptide or Mn^2+^_N_ in the structure, indicating that crystals may not be stable at this pH (Table S3).

We then utilized quantum chemistry calculations to assess the effect of the protonation state of His333 on the local geometry around Mn^2+^_N_. The geometry-optimized structure with neutral His333 depicts a strongly bonded to Mn^2+^_N_ as indicated by the bond distance of ∼2.2 Å; Glu554 is also H-bonded to two hydroxyls of GalNAc (Fig. 3b, middle panel). At an artificially low pH with 100% [His333] ^+^, such strong interactions are not seen (Fig. 3b, right panel). His333 does not coordinate Mn^2+^_N_ and Glu554 H-bonds with one hydroxyl of GalNAc, suggesting weaker binding to the glycopeptide substrate (Fig. 3b, right panel). The difference in Mn^2+^_N_ and glycopeptide binding at varying pHs may explain why the peptide ligands dissociate easily from the binding pocket when His333 is protonated. At pH 7, His is at an equilibrium between uncharged (∼90%) and charged (∼10%) species. Thus, the E:S complex can readily form when His333 is uncharged and coordinating Mn^2+^_N_, catalysis can occur, and the product is released due to His333 protonation (Fig. 3b, middle panel). Changing His333 to Asn decouples Mn^2+^_B_ and GalNAc binding and results in a ∼30-fold decrease in the K_M_ and ∼125-fold decrease in V_max_ compared to *T.gondii*-GalNAc-T3^WT^, further showing that Mn^2+^_N_ coordination is fine-tuning substrate binding and catalysis in a pH-dependent manner (Fig. S4b and S5d, Table S2).

### An active site loop (II) modulates substrate binding

Metazoan GalNAc-Ts contain a catalytic flexible gating loop (loop I) that becomes ordered upon UDP-GalNAc binding and has an additional role in peptide binding ^20,21,24^. Loop I is conserved in *T.gondii*-GalNAc-T3 (His414-Pro426, Fig. 4a). GalNAc-Ts contain an additional loop (Loop II) in the active site (Fig. 4b), and in *T.gondii*-GalNAc-T3 and its Apicomplexan homologues, it mainly consists of small hydrophobic residues (Ala317-Cys322) including Gly319, which makes a mainchain interaction with the peptide backbone (Fig. 4a). Loop II is disordered in the apo structure and becomes more ordered upon peptide substrate binding (Fig. 4a). Loop II flexibility and the interaction that occurs between the Gly319 backbone carbonyl and an amide group on both di-glycopeptide substrates (Muc5AC-3,13 and Cst1.4) allow *T.gondii*-GalNAc-T3 to accommodate and modify substrates with variable sequences. Substituting the central loop residue Ile320 with Pro to perturb the conformation and stability of the loop diminishes enzymatic activity, supporting a possible role for loop II in substrate binding and alignment (Fig. 4c).

**Fig. 4:**
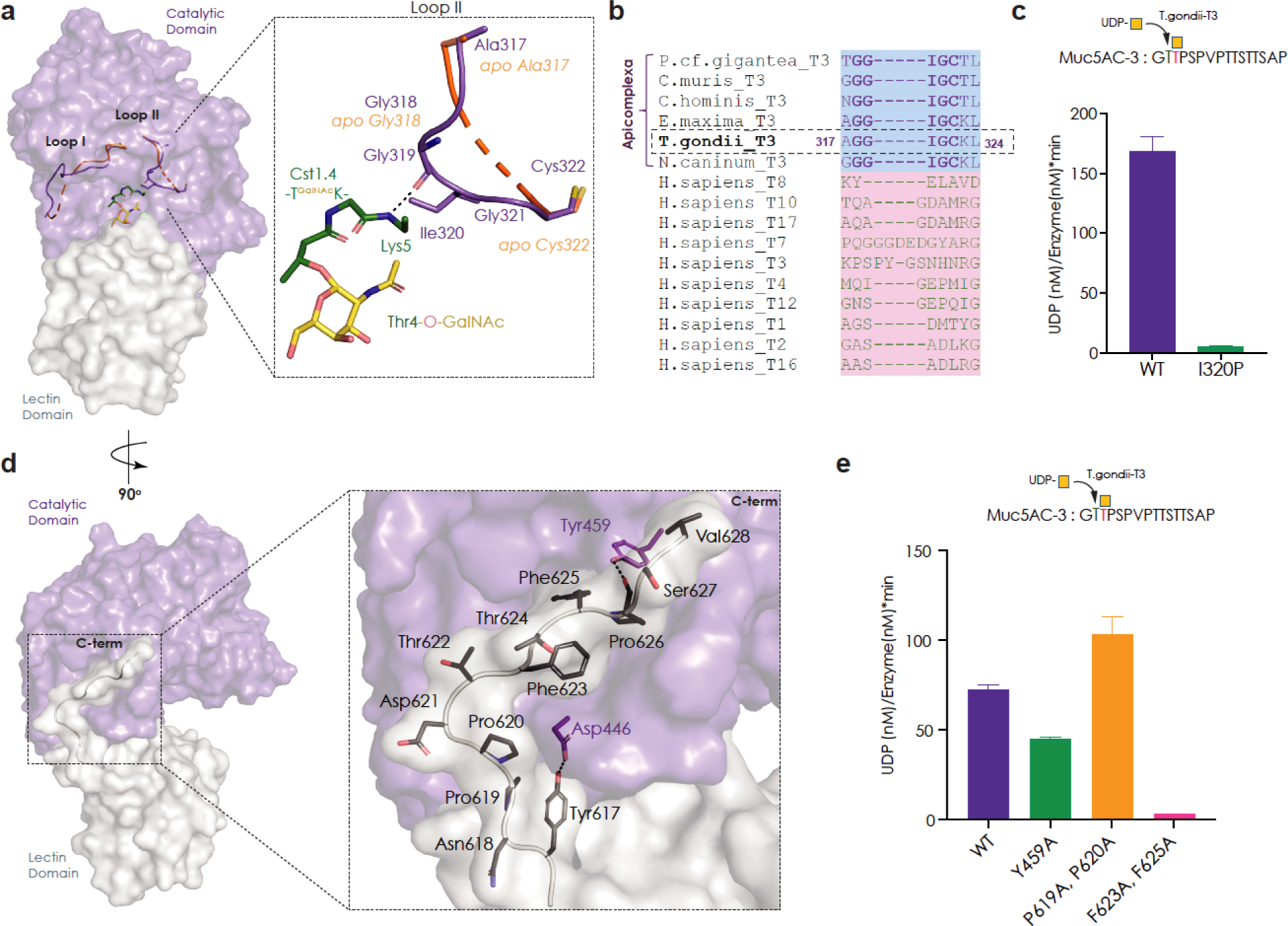
A second substrate binding loop (II) and an extended C-terminal tail influence enzymatic function. **a,** GalNAc-Ts contain a catalytic gating loop that helps align UDP-GalNAc in the active site (Loop I). *T.gondii*-GalNAc-T3 contains an additional loop (loop II) that interacts with the peptide backbone **b,** A sequence alignment comparing loop II in *T.gondii*-GalNAc-T3 to Apicomplexan homologues and human GalNAc-Ts. Loop II is unique to *T.gondii*-GalNAc-T3 and its homologues. **c,** Inserting a Pro in the middle of the loop (I320P) disrupts enzymatic activity. **d,** *T.gondii*-GalNAc-T3 contains an extended C-terminal tail that makes extensive hydrophobic and H-bonding interactions with the catalytic domain. **e,** P619A/P620A double mutation decreases loop rigidity to slightly increase enzymatic function. Reducing loop hydrophobicity (F623A/F625A) disrupts enzymatic function.

### An extended hydrophobic C-terminal tail stabilizes the conformation of *T.gondii*-GalNAc-T3

*T.gondii*-GalNAc-T3 contains a non-conserved extended C-terminal tail (Tyr617-Val628) that interacts with the catalytic domain via hydrophobic and H-bonding interactions (Fig. 4d). Given the extensive nature of these interactions, we hypothesize that the extended C-terminus stabilizes the enzyme conformation and restricts the movement of the lectin domain. Tyr617 of the C-terminal tail H-bonds with Asp446 in the catalytic domain to hold the N-terminal end of loop in position, while residues Pro619-Pro620 behave like rigid stand before the tail bends to position Phe623 and Phe625 into a hydrophobic cleft of the catalytic domain. Finally, an H-bond between Tyr459 in the catalytic domain with the mainchain of Pro626 holds down the C-terminal end of loop. Attempts to mutate Tyr617 and truncate the C-terminal tail abrogated protein expression in yeast, suggesting destabilization of *T.gondii*-GalNAc-T3. We observe a ∼40% reduction in activity for Y459A and significantly reduced activity for F623A/F625A (Fig. 4e). Interestingly, increasing the flexibility of the tail in the P619A/P620A variant results in a ∼25% increase in activity compared to *T.gondii*-GalNAc-T3^WT^.

### The *T.gondii*-GalNAc-T3 lectin domain does not influence activity

The lectin domain consists of 3 repeats (α, β, and γ) each with the potential to interact with GalNAc if the residues Asp-His-Asn are present (Fig. S6a). In certain metazoan GalNAc-Ts, lectin domain interactions with a Thr-O-GalNAc ∼9-12 amino acids away from the acceptor has long-range enhancing effects on activity. Although none of the *T.gondii*-GalNAc-T3 lectin repeats contain the Asp-His-Asn GalNAc binding motif, we wondered if *T.gondii*-GalNAc-T3 uses a distinct mode of GalNAc recognition to enhance activity. However, we do not observe enhancement in activity when comparing *T.gondii*-GalNAc-T3 activity towards di-glycopeptides that contain one GalNAc at the +1 position and another GalNAc at either +12 (DGPI) or −10 (DGPII) to mono-glycopeptides with GalNAc only at the +1 position (GPIID), and thus have no evidence for a role for the lectin domain in activity enhancement by interacting with a distal GalNAc on a di-glycopeptide (Fig. S6b).

### Mn^2+^_N_ site mutations and C-terminal tail truncations disrupt *T. gondii* bradyzoite cyst wall O-glycosylation

To assess the significance of the *in vitro* characterizations of *T.gondii*-GalNAc-T3 in an intact organism, *T. gondii* cell lines expressing various Mn^2+^_N_ and C-terminal tail mutants were generated by genetically manipulating the T3 locus. Mutations in residues constituting the Mn^2+^_N_ site and a C-terminal tail truncation resulted in the disruption of cyst wall O-glycosylation as detected by a glycoepitope-specific antibody (Fig. 5). Interestingly, although *T.gondii*-GalNAc-T3^H333A^ results in diminished activity *in vitro* (Fig. 2d), it does not completely abolish cyst wall O-glycosylation *in vivo*, suggesting the presence of additional co-factors or conditions that influence *T.gondii*-GalNAc-T3 function *in vivo*.

**Fig. 5:**
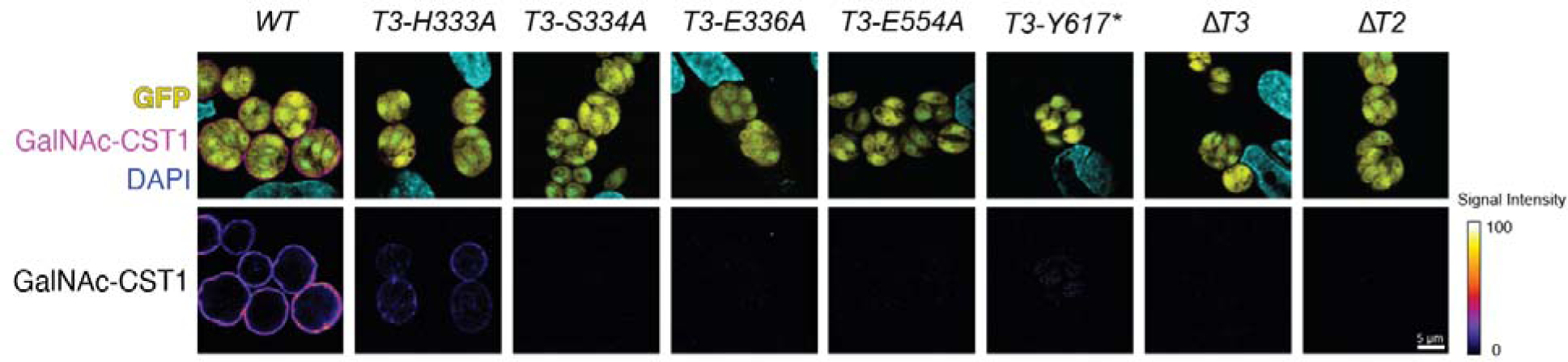
The second metal site and C-terminal tail influence in vivo O-glycosylation. *In vitro T. gondii* cysts in HFF cells probed with GalNAc glyco-epitope-specific anti-CST1 mucin antibody reveal that the glycosylation of the cyst wall is diminished in *T.gondii*-GalNAc-T3 mutant parasites. GFP expression indicates that the parasites are differentiated into cyst-forming bradyzoites. The lower panels delineate the reduced intensity of cyst wall glycosylation.

To evaluate the extent of glycosylation within the mucin domain of CST1, we introduced a surrogate protein comprising the CST1 signal peptide, CST1 mucin domain, and hemagglutinin (HA) epitope tag without SRS domains. The protein was expressed in both WT and various *T.gondii*-GalNAc-T3 mutants and the resulting parasite lysates were analyzed by immunoblotting of the HA tag (Fig. S7). In *T.gondii*-GalNAc-T3^WT^ cells, the mucin domain is fully O-glycosylated, as evidenced by the migration pattern of the surrogate mucin, which remained at the interface between the stacking and resolving gel. In contrast, the H333A mutant exhibited intermediately glycosylated species with molecular weight around ∼200 kDa. Furthermore, both lower molecular weight species at 55 kDa and fully glycosylated forms are detected, suggesting that *T.gondii*-GalNAc-T3^H333A^ retains partial enzymatic activity *in vivo*. As expected, the remaining mutants affecting the Mn^2+^_N_ binding site completely abolished *T.gondii*-GalNAc-T3 enzymatic activity, as indicated by the presence of lower molecular weight species in the immunoblots. In the case of the C-terminal truncation mutant, only intermediately glycosylated species were generated, with fully glycosylated forms conspicuously absent. In summary, mutations in the second metal binding site and C-terminal region of T.gondii-GalNAc-T3 disrupt bradyzoite cyst wall glycosylation, with varying effects on enzymatic activity and mucin domain glycosylation.

## Discussion

We report the first structures of a protozoan GalNAc-T, *T.gondii*-GalNAc-T3, and provide mechanistic insights into novel druggable sites to treat latent toxoplasmosis and other related parasitic diseases that are currently challenging to manage. In addition, we fully characterize the *T.gondii*-GalNAc-T3 substrate preference for GalNAc at the +1 position of an acceptor on substrate using a broad repertoire of (glyco)peptides and show that the transferase can fully glycosylate a stretch of Thr *in vitro* by a processive mechanism. This preference is shared by human GalNAc-T10, T7, and T17, but these isoenzymes do not contain a similar motif and most likely use a distinct substrate binding pocket to interact with Thr-O-GalNAc. The structures reveal novel characteristics that are critical for the function of *T.gondii*-GalNAc-T3 *in vitro* and *in vivo* and strictly conserved among its Apicomplexan homologues, including a unique 2^nd^ metal (Mn^2+^_N_), a novel active site residue (Glu332), a dynamic substrate binding loop (II), and an extended C-terminus (Fig. 6a).

**Fig. 6:**
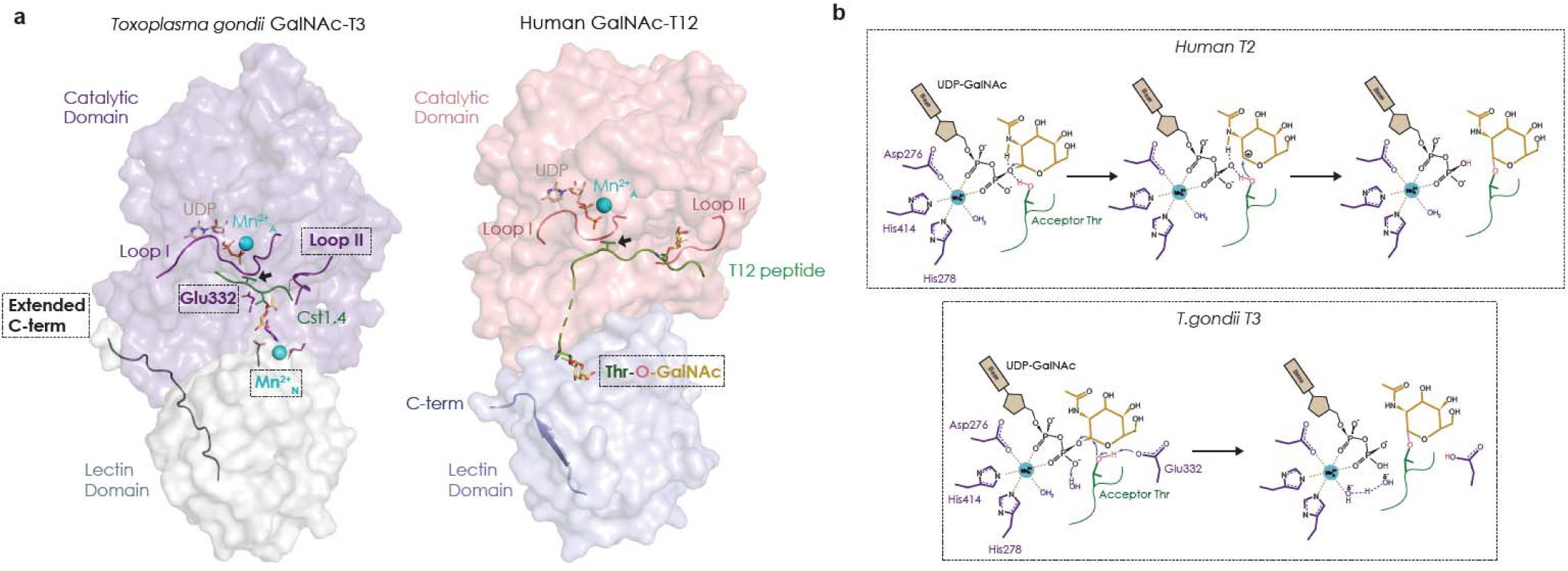
Catalytic mechanism of *T.gondii*-GalNAc-T3. **a,** A comparison of *T.gondii*-GalNAc-T3 structure to human GalNAc-T12 reveals distinct features that could be targets for Toxoplasmosis therapeutics. *T.gondii*-GalNAc-T3 has an extended C-terminal tail, a second Mn^2+^_N_, a flexible loop II that interacts with substrate peptide mainchain, and a putative general base Glu332 in the active site, while human GalNAc-T12 contains a GalNAc binding pocket in the lectin domain, which does not seem to be present in the lectin domain of *T.gondii*-GalNAc-T3. **b, top panel,** Catalytic mechanism of human GalNAc-T2. The acceptor Thr approaches the anomeric carbon in a front face reaction, resulting in the formation of an oxocarbenium ion. The pyrophosphate leaving group extracts a proton from Thr as a bond forms between the acceptor and GalNAc. **b, bottom panel,** Proposed catalytic mechanism of *T.gondii*-GalNAc-T3. Glu332 could act as a general base and extract a proton from the acceptor Thr, while it forms a bond with the anomeric carbon of GalNAc in an S_N_2 type nucleophilic reaction. In this case, the leaving pyrophosphate group could extract a proton from a nearby water in the active site.

Mn^2+^_N_ is coupled to the GalNAc binding pocket that dictates the *T.gondii*-GalNAc-T3 +1-substrate preference. Mn^2+^_N_ coordination by His333 and Glu554 does not align these sidechains for GalNAc binding, but instead, appears to influence GalNAc binding by altering their charges (Fig. 3a). The second metal site is conceivably sensitive to metal concentrations *in vivo*, where changes in metal concentration during parasite invasion and replication could regulate glycosylation. While little is known about the role of Mn^2+^ in *T. gondii*, Ca^2+^ is critical for *T. gondii* function, and uptake of extracellular Ca^2+^ modulates host invasion ^29^. Thus, it is possible that *T.gondii*-GalNAc-T3 function is fine-tuned by Ca^2+^ binding to the second metal site during host infection to glycosylate virulence factors. Since Ca^2+^ has less stringent metal coordination rules than Mn^2+^, it may bind more efficiently in the absence of His333, which could explain why we see more glycosylation *in vivo* than *in vitro* at physiological concentrations of Ca^2+^, which is the subject of current studies on the enzyme. Other unknown factors could also contribute to the *in vivo* activity of *T.gondii*-GalNAc-T3^H333A^.

Our structures revealed an unexpected active site residue Glu322, which influences reaction chemistry. The current catalytic mechanism of human GalNAc-T2 is proposed to be S_N_1 or S_N_i type^30^, where the acceptor Thr hydroxyl uses a front-face reaction to form a bond with the anomeric carbon of the GalNAc moiety of the donor substrate. In the first step of catalysis, protons on the leaving group UDP phosphate and amide group of GalNAc stabilize the positive charge that develops on the anomeric carbon to form an oxocarbenium ion as the acceptor Thr approaches (Fig. 6b, top panel). The reaction does not use a general base, and transfer of the hydroxyl hydrogen to the phosphate leaving group occurs as the bond between GalNAc and Thr forms. In *T.gondii*-GalNAc-T3, the acceptor Thr is similarly positioned for front face attack. However, the orientation and distance of Glu322 to the acceptor hydroxyl indicates that it could act as a general base to deprotonate the acceptor Thr prior to nucleophilic attack in an S_N_2-type reaction (Fig. 6b, bottom panel). In this scenario, the leaving group phosphate extracts a proton from another source, such as the nearby water molecule. It is possible that the second metal has long range effects on the electrostatic environment of the active site that requires the presence of Glu332 to compensate for these effects that are not present in metazoan homologues. We are currently conducting computational and structural studies to investigate the catalytic mechanism of *T.gondii*-GalNAc-T3.

The other differences between *T.gondii*-GalNAc-T3 and its metazoan homologues include the ordering of Loop II due to non-specific backbone interactions substrates, helping to align various substrates across the active site, while the C-terminal tail stabilizes an active conformation of the enzyme. Loop II is more variable in metazoan GalNAc-Ts, and in most crystal structures is not contacting the substrate (Fig. 4b). The exceptions are human GalNAc-T2 in complex with an unglycosylated peptide EA2, where loop II makes mainchain interactions with the peptide backbone, and GalNAc-T12, where the sidechain of Asn270 in loop II forms part of the GalNAc binding pocket and enhances glycopeptide substrate binding^20,24^. Whether these residues are required for substrate binding in human GalNAc-T2 is not clear, but the loop is ordered and not contacting the substrate in complex structures of GalNAc-T2 with glycopeptides, suggesting it may enhance peptide, but not glycopeptide binding.

Intriguingly, the lectin domain does not enhance activity by binding to distant sites, as seen with metazoan enzymes, but instead contains Glu554, which binds to Mn^2+^_N_ and positions the C-terminal tail in the catalytic domain cleft. The possible inability of the *T.gondii*-GalNAc-T3 lectin domain to enhance activity by binding GalNAc is consistent with the sequential mechanism it uses to efficiently glycosylate stretches of Thr without needing to bind a long range GalNAc. Whether the lectin domain has additional functions in *T.gondii*-GalNAc-T3 such as in binding substrates or co-factors remains to be studied.

There are currently no treatments that can eliminate *T. gondii* tissue cysts and new medications to eradicate latent toxoplasmosis are clearly needed. The differences between *T.gondii*-GalNAc-T3 and human GalNAc-Ts represent regions that can be potentially targeted for therapeutic purposes, an area we are currently exploring (Fig. 6a). Given the similarities between *T.gondii*-GalNAc-T3 and its Apicomplexan homologues, it is conceivable that inhibitors of *T.gondii*-GalNAc-T3 could be used more widely to target other disease-causing parasites, such as *Cryptosporidium hominis* which causes the gastrointestinal infection cryptosporidiosis, as well as *Neospora caninum* and *Eimeria maxima*, which infect livestock resulting in agricultural economic losses. In conclusion, our studies of *T.gondii*-GalNAc-T3 have shed light on the evolution of this enzyme family and lay the groundwork for future studies on anti-microbials that target toxoplasmosis and other parasitic diseases.

## Data availability

Structure coordinates and X-ray diffraction data have been deposited in the Protein Data Bank, www.wwpdb.org (PDB ID codes:). All data are available in the main text or the supplementary materials.

## Acknowledgements

We thank the beamline staff at the Advanced Photon Source for assistance. The quantum chemical study utilized the computational resources of the NIH HPC Biowulf cluster (http://hpc.nih.gov). We thank our colleagues Dr. Wei Yang, Dr. Kelly Ten Hagen, and Dr. Lawrence Tabak for reading the manuscript and providing feedback. This work was supported by National Institutes of Health Grant 1-ZIA-DE000754-03 (to N.L.S.) and 5-R01-GM113534-08 (to T.A.G.).

## Author information

### Contributions

P.K. and N.L.S. conceptualized the project. P.K. cloned, expressed, and purified *T.gondii*-GalNAc-T3. P.K. crystallized *T.gondii*-GalNAc-T3 and co-complexes and conducted *in vitro* biochemical assays. N.L.S. and P.K. harvested crystals, and collected data, and P.K. processed the data and solved and refined the structures. T.A.G. conducted ^3^H-GalNAc assays on glycopeptide substrates and glycosylation site analysis by Edman sequencing and C.J.B. performed kinetic studies on the Ser and Thr glycopeptides. T.T. and L.M.W. performed the *in vivo* O-glycosylation studies. N.L.S. prepared the manuscript. All authors contributed to the results and methods section and in editing the manuscript.

## Methods

### Expression and purification of *T.gondii*-GalNAc-T3

*T.gondii*-GalNAc-T3 (aa 74–635) was cloned from a *T.gondii*-GalNAc-T3 cDNA template (GenScript AY160970.1, GeneID:7901231, TGME49_318730) into the expression vector pPICZα-A (Invitrogen) for secreted protein expression in *Pichia pastoris*. *T.gondii*-GalNAc-T3 mutant constructs were made by site-directed mutagenesis (Table S4). For strain construction, cloned plasmids were linearized with the restriction enzyme PmeI and transformed into *Pichia pastoris* SMD1168 cells (Invitrogen) by electroporation. To express *T.gondii*-GalNAc-T3, cells were grown at 30°C to an OD600 ∼ 20 in BMGY media (2 % (w/v) peptone, 1 % (w/v) yeast extract, 1.34 % (w/v) Yeast Nitrogen Base (YNB), 4 × 10^-5^ % (w/v) biotin, 1 % (v/v) glycerol, 100 mM potassium phosphate pH 6.0) containing 100 μg/ml of Zeocin (Invivogen). To induce protein expression, cells were cultured by centrifugation (1500 X g for 10 min) and resuspended in 1/5 volume BMMY media (1% glycerol is replaced with 0.5% methanol) and incubated at 20°C for 24 h.

Cells were centrifuged (1500 X g for 10 min) and supernatant was collected, filtered, and pH adjusted by adding 50 mM Tris pH 8.0, 250 mM NaCl, 5% glycerol and 10 mM β-mercaptoethanol (βME). Purification was carried out at 4°C and supernatant was applied onto a 5 ml HisTrap HP column (Cytiva) pre-equilibrated with 5 column volumes (CV) of buffer A (25 mM Tris, 250 mM NaCl, 10 mM βME and 5% glycerol, pH 7.5). Protein was eluted by a 50-500 mM imidazole gradient over 10 CV. Peak fractions were pooled and incubated with TEV protease at a ratio of 1:20 (w/w) while dialyzing into 500 ml of buffer A containing 25 mM imidazole at 4°C. The His-tagged TEV protease and residual uncut protein were removed by manually loading the sample onto a 1 ml HisTrap HP column (Cytiva) equilibrated with 5 CV dialysis buffer, followed by washing with 1 CV of dialysis buffer. After increasing the buffer glycerol concentration to 30 %, protein was aliquoted, flash frozen in LN2, and stored at −80°C.

### Probing neighboring glycosylation activity against glycopeptide GPIIC

Assays against random glycopeptide GPIIC (Fig 1b) consisted of 100 μl reactions consisting of 100 mM sodium cacodylic pH 6.5, 0.8 mM β-mercaptoethanol, 0.08% Triton X100, 10 mM Mn^2+^, ∼0.8 mM GPIIC glycopeptide, 0.4 mM UDP-[³H]GalNAc, 1.4 μM *T.gondii*-GalNAc-T3, were incubated at 37°C on a shaking microincubator for 5 hr and 21 hr. Reactions were quenched with 100 μl of 250 mM EDTA, diluted to 4 ml and passed over a 1 ml column of DOWEX 1X8 (Sigma Aldrich). Dowex column flow through was lyophilized and applied to a Sephadex G10 column (Cytiva) to separate peptide from free [³H]GalNAc and the peptide peak was lyophilized for sequencing. Glycopeptide products were Edman sequenced on a modified gradient Shimadzu PPSQ53A sequencer (Shimadzu Scientific instruments Inc., Columbia, MD). The ^3^H-GalNAc-O-Thr-PTH derivatives (11-14.5 min) at each cycle were collected on a Shimadzu FRC-10A fraction collector and ^3^H-scintillation counted on a Beckman LS650 scintillation counter.

### Probing lectin domain long range glycosylation activity against di-glycopeptides

To probe lectin domain interactions, 100 μl reactions consisting of 100 mM sodium cacodylic pH 6.5, 0.8 mM β-mercaptoethanol, 0.08 % Triton X100, 10 mM Mn^2+^, ∼1.7 mM (glyco)peptides DGPI, DGPII and GPIID (Supplemental figure S6b), 0.1 mM UDP-[³H]GalNAc, 0.7 μM *T.gondii*-GalNAc-T3, incubated at 37°C for 5 hr. Reactions were quenched with 100 μl 250 mM EDTA and processed as described for GPIIC. Pre and post Dowex ^3^H DPM and ^3^H DPM integration of the Sephadex G10 glycopeptide and GalNAc peaks were used to calculate percent of glycopeptide glycosylated and percent of UDP-[³H]GalNAc hydrolysis.

### Identification of sequential glycosylation sites in *T. gondii* peptides and Muc5AC by Edman sequencing

For peptide O-glycosylation analysis by Edman Sequencing, 41-46 μl reactions consisting of 110-120 mM sodium cacodylic pH 6.5, 0.8 mM β-mercaptoethanol, 0.08% Triton X100, 12 mM Mn^2+^, 0.5-0.6 mM (glyco)peptides, 2.2-2.4 mM UDP-[³H]GalNAc, 0.06-0.15 μM *T.gondii*-GalNAc-T3, incubated at 37°C for 30 min (Cst1.x) or 90 min (Srs3.x & Muc5ACx) (Fig 1d and Supplemental figure S1c,d,e). Overnight incubations were also done for Cst1.3 and Muc5AC-13 (Fig 1d, Supplemental figure S1d,g). Reactions were quenched with 100 μl 250 mM EDTA and processed as described for GPIIC. Pre and post Dowex ^3^H DPM and ^3^H DPM integration of the Sephadex G10 glycopeptide and GalNAc peaks were used to calculate percent of glycopeptide glycosylated and percent of UDP-[³H]GalNAc hydrolysis. Glycopeptide products were Edman sequenced on a modified gradient Shimadzu PPSQ53A sequencer. Free GalNAc (2.25-3.25 min) and the GalNAc-O-Thr-PTH and GalNAc-O-Ser-PTH derivatives (9.0-14.5 min) at each cycle were collected on a Shimadzu FRC-10A fraction collector and each fraction scintillation counted on a Beckman LS650 scintillation counter. Sequence chromatograms of the PTH (phenylthiohydantion) derivatives were also analyzed as described in Supplemental figure S1g.

### Kinetics of *T.gondii*-GalNAc-T3 against Ser-O-GalNAc and Thr-O-GalNAc glycopeptides

Stock solutions of glycopeptide substrates T7T* and T7S* (Supplemental figure S1f) were made to yield final reaction concentrations of 1.4 mM, 0.7 mM, 0.35 mM, 0.175 mM, and 0.88 mM. Reactions consisted of 100 mM sodium cacodylic pH 6.5, 1 mM β-mercaptoethanol, 0.1% Triton X-100, 2 mM UDP-[³H]GalNAc, 0.044 µM enzyme, peptide substrate, and were incubated at 37°C. Reaction times varied, depending on substrate concentrations, ranging from 10 to 30 min to maintain peptide glycosylation to <20%. After incubation, reactions were quenched with 200 µL of 0.5% TFA in H_2_O. BioPureSPIN TARGA-C18 spin columns (The Nest Group, Ipswitch MA) were prior equilibrated by passing sequentially: acetonitrile (300 µL), 50/50 acetonitrile/H_2_O in 0.1% TFA (300 µL), and 0.1% TFA in H_2_O (700 µL). The latter washes were eluted by spinning at 800rpm in an Eppendorf Minispin Plus tabletop centrifuge. Ten percent (22 µL) of the reaction volume was removed for [3H] scintillation counting (initial DPM), and the remainder was applied to the equilibrated TARGA C18 hydrophobic spin columns and spun for 1 minute. After the sample was eluted columns were washed with 800 µL of 0.1% TFA in H_2_O to remove free UDP-[³H]GalNAc and [³H]GalNAc by centrifugation at 800 rpm for 1 min giving the A-eluate. The bound (glyco)peptide products/reactants (B eluate) were eluted using two washes of 200 µL of 50/50 acetonitrile in 0.1% TFA followed by 200 µL of 100% acetonitrile, each spun for 1 min at 800 rpm, and a final 100 µL of 100% acetonitrile spun for 4 min. [^3^H] scintillation counting was performed on the combined flow through and wash (A eluate) and the eluted (glyco)peptide products/reactants (B eluate). The extent of peptide glycosylated, in mM, was obtained by dividing the B counts (in DPM) of the eluted (glyco)peptides by the initial total DPM (as well as by the sum of the DPM of the A and B eluates) of the UDP-[³H]GalNAc and by multiplying by the initial mM of UDP-GalNAc. Values were converted to µM of GalNAc transferred/(µM enzyme*min) according to the initial amount of UDP-GalNAc, substrate, and enzyme used. K_cat_ (µM GalNAc/(µM enzyme*min) or min^-1^), K_M_ (µM), and V_max_ (µM GalNAc/min) values and Michaelis Menten plots were obtained using GraphPad Prism software (Boston, MA).

### Crystallization, data collection and processing, structure determination and refinement

*T.gondii*-GalNAc-T3 was thawed and exchanged into crystallization buffer (25 mM HEPES, 100 mM NaCl, 0.5 mM EDTA, 5% Glycerol and 10 mM βME, pH 7.3) on a Superdex 16/600 HiLoad column (Cytiva). Peak fractions were concentrated using a 10 kDa cut-off Amicon ultra-concentrator (Millipore Sigma) to ∼10-15 mg/ml. Each enzyme-peptide-UDP-Mn^2+^ complex was prepared by combining *T.gondii*-GalNAc-T3, 5 mM of one of the glycopeptides (Muc5AC-3, Muc5AC-13, Muc5AC-3,13, Cst1.4, and Srs13.2 (Anaspec, Fremont, CA), 5 mM UDP-2-(acetylamino)-4-F-D-galactosamine disodium salt (UDP-GalNAc-F), and 5 mM MnCl_2_ to a final protein concentration of 6.0 mg/ml. Hanging drops were prepared by mixing 1 μl of protein complex solution with 1 μl of reservoir solution containing 0.1 M CHES pH 9.5 and 14-20% PEG 8000 (w/v) and equilibrated against a 500 μl reservoir solution. Crystals formed after 4 days in 24-well plates incubated at 20°C. Crystals were cryoprotected in a crystallization solution containing 20% glycerol and flash frozen in LN2 prior to X-ray data collection.

X-ray data was collected at the Advanced Photon Source SER-CAT ID and BM-22 beam lines (Argonne, IL). HKL2000 was used to process and scale the X-ray diffraction data (Tables S2 and S3)^31^. The initial structure was solved by molecular replacement using MolRep (CCP4) and an Alphafold2 model of *T.gondii*-GalNAc-T3 as an initial search model^32–34^. Initial models were rebuilt manually using Coot and refined in PHENIX^35,36^. The final models were validated by using PROCHECK and MOLPROBITY^37–40^. Structure figures were prepared with Pymol (The PyMOL Molecular Graphics System, Version 2.0 Schrodinger, LLC).

### Quantum Chemistry

From the crystal structure of *T.gondii*-GalNAc-T3 in complex with a glycopeptide substrate, we chose His333, Ser334, Tyr335, Glu336, Glu554 and a portion of glycopeptide substrate as well as the Mn^2+^ and water molecules for quantum chemical calculations at the level of density functional theory. Their backbone and side chains were then modified; see the coordinates given in the Supporting info. Quantum chemical calculations were carried out with Gaussian 16^41^ on the neutral and protonated His333. We employed M06-L with the basis set of cc-PVDZ in the water reaction field for geometry optimization.

### UDP-Glo enzymatic assay

Glycosyltransferase activity was assayed using a UDP-Glo^TM^ Glycosyltransferase Assay kit (Promega, V6961). A 25 μL reaction was initiated by adding 50 nM and 100 nM of purified wild type and mutant *T.gondii*-GalNAc-T3, respectively, to a 5 mM donor substrate uridine diphosphate (UDP)-N-acetylgalactosamine (GalNAc) (supplied within the kit) and acceptor substrate; Muc5AC, Muc5AC-3, Muc5AC-13, Muc5AC-3,13, Cst1.1, Cst1.2, Cst1.3, Cst1.4, Srs13.1, Srs13.2, Srs13.3, and Srs13.4 peptides and glycopeptides (Anaspec) in 5 mM MnCl_2_ (Sigma Aldrich) and 1 X HEPES buffer (100 mM NaCl, 5 mM βME, 25 mM HEPES, pH 7.3). Reactions were assembled in 96-well plate (Corning) and incubated at 37°C for 15 min. The reaction was terminated by adding 25 μL UDP detection reagent to each well followed by incubation at 27°C for 60 min in BioTak plate reader, and Luminescence was recorded. A standard curve of UDP in 5 mM MnCl_2_ and 1X HEPES buffer was applied at each measurement to associate a defined UDP concentration with a luminescence signal that directly correlates with the glycosyltransferase activity within 60 min under described conditions. Multiple reactions with varying substrate concentrations (0-2000 μM) were used to determine the kinetic parameters of the glycosyltransferase reactions. All reactions were performed in triplicate. Data were analyzed by using Microsoft Excel and GraphPad Prism software (Boston, MA).

### *T. gondii* cell culture and strains

Prugniaud strain with a deletion in KU80 gene^42^ was cultured in human foreskin fibroblasts (HFF) in 10% fetal bovine serum (FBS) in Dulbecco’s modified Eagle medium (DMEM) with penicillin-streptomycin at 5% CO_2_. For the induction of bradyzoite differentiation, DMEM with 1% FBS with 25 mM HEPES adjusted to pH 8.2 was used at atmospheric CO_2_.

### Genetic manipulation of *T. gondii*

For generating point mutations on *T.gondii*-GalNAc-T3 gene in *T. gondii*, gRNAs targeting the *T.gondii*-GalNAc-T3 locus was used with the donor oligos that repair with the desired point mutation as described previously^7,43^. Briefly, two candidate gRNA sequences were selected for three loci (His333/Ser334/Glu336, Glu554, and Tyr617). The gRNA oligos were incorporated into gRNA-Cas9 vector using NEBuilder HiFi DNA Assembly kit (New England Biolabs). Each donor oligo contains a point mutation that replace original codon with alanine or stop codon flanked by 40 nt homologous recombination sequences on both 5’ and 3’ (all donor sequences and gRNA vectors are shown in supplementary files). The gRNA-Cas9 vectors and corresponding donor oligos were electroporated into the parental Pru strains and subcloned by limiting dilution. The point mutations were verified by Sanger sequencing the genomic DNA of the parasite clones. Surrogate mucin construct was generated by NEBuilder by concatenating the constitutive promoter, CST1 signal peptide, CST1 mucin domain, 3x HA sequences, and selectable marker DHFR. The surrogate mucin construct was electroporated into the parasite and integrated into genome by pyrimethamine selection.

### Immunofluorescence assay and immunoblotting

HFF cells, grown on a coverslip infected with *T.gondii*-GalNAc-T3 point mutant *T. gondii*, were cultured in a bradyzoite differentiation medium for 72 hours, as previously described. Following incubation, cells were fixed with 4% paraformaldehyde in PBS for 30 minutes and subsequently permeabilized with 0.2% Triton-X100 in PBS for 20 minutes. For immunostaining, cells were incubated with a 1:200 dilution of rabbit anti-GFP antibody (ThermoFisher #G10362) and GalNAc glycoepitope-specific anti-CST1 antibody^6^, both prepared in PBS with 1% BSA. The incubation was carried out for 90 minutes at 37°C. Secondary fluorescent antibodies were applied at a 1:2000 dilution in PBS with 1% BSA for another 90 minutes. For immunoblot analyses, HFF cells infected with *T. gondii* strains expressing the surrogate mucin protein with an HA tag were cultured for 48 hours in standard medium. Cells were harvested, lysed in Laemmli SDS buffer, and the lysates subjected to SDS-PAGE. Following transfer, the blots were probed with HRP-conjugated rat anti-HA antibody (clone 3F10, Roche) at a 1:1000 dilution and incubated overnight. Signal detection was accomplished using the SuperSignal West Pico Plus reagent (ThermoFisher).

## References

1 Montoya, J. G. & Liesenfeld, O. Toxoplasmosis. Lancet 363, 1965–1976 (2004). 10.1016/S0140-6736(04)16412-X

2 Schluter, D. et al. Animals are key to human toxoplasmosis. Int J Med Microbiol 304, 917–929 (2014). 10.1016/j.ijmm.2014.09.002

3 Ford, N. et al. Managing Advanced HIV Disease in a Public Health Approach. Clin Infect Dis 66, S106-SS110 (2018). 10.1093/cid/cix1139

4 Alday, P. H. & Doggett, J. S. Drugs in development for toxoplasmosis: advances, challenges, and current status. Drug Des Devel Ther 11, 273–293 (2017). 10.2147/DDDT.S60973

5 Wandall, H. H., Nielsen, M. A. I., King-Smith, S., de Haan, N. & Bagdonaite, I. Global functions of O-glycosylation: promises and challenges in O-glycobiology. FEBS J 288, 7183–7212 (2021). 10.1111/febs.16148

6 Tomita, T. et al. The Toxoplasma gondii cyst wall protein CST1 is critical for cyst wall integrity and promotes bradyzoite persistence. PLoS Pathog 9, e1003823 (2013). 10.1371/journal.ppat.1003823

7 Tomita, T., Ma, Y. & Weiss, L. Characterization of a SRS13: a new cyst wall mucin-like domain containing protein. Parasitol Res 117, 2457–2466 (2018). 10.1007/s00436-018-5934-3

8 Craver, M. P., Rooney, P. J. & Knoll, L. J. Isolation of Toxoplasma gondii development mutants identifies a potential proteophosphogylcan that enhances cyst wall formation. Mol Biochem Parasitol 169, 120–123 (2010). 10.1016/j.molbiopara.2009.10.006

9 Zinecker, C., Striepen, B., Tomavo, S., Dubremetz, J. & Schwartz, R. T. The dense granule antigen, GRA2 of Toxoplasma gondii is a glycoprotein containing O-linked oligosaccharides. Molecular and Biochemical Parasitology 97, 241–246 (1998).

10 Tomita, T. et al. Making Home Sweet and Sturdy: Toxoplasma gondii ppGalNAc-Ts Glycosylate in Hierarchical Order and Confer Cyst Wall Rigidity. mBio 8 (2017). 10.1128/mBio.02048-16

11 Raman, J., Guan, Y., Perrine, C. L., Gerken, T. A. & Tabak, L. A. UDP-N-acetyl-alpha-D-galactosamine:polypeptide N-acetylgalactosaminyltransferases: completion of the family tree. Glycobiology 22, 768–777 (2012). 10.1093/glycob/cwr183

12 Hazes, B. The (QxW)3 domain: a flexible lectin scaffold. Protein Sci 5, 1490–1501 (1996). 10.1002/pro.5560050805

13 Wojczyk, B. S. et al. cDNA cloning and expression of UDP-N-acetyl-D-galactosamine:polypeptide N-acetylgalactosaminyltransferase T1 from Toxoplasma gondii. Mol Biochem Parasitol 131, 93–107 (2003). 10.1016/s0166-6851(03)00196-8

14 Stwora-Wojczyk, M. M., Dzierszinski, F., Roos, D. S., Spitalnik, S. L. & Wojczyk, B. S. Functional characterization of a novel Toxoplasma gondii glycosyltransferase: UDP-N-acetyl-D-galactosamine:polypeptide N-acetylgalactosaminyltransferase-T3. Arch Biochem Biophys 426, 231–240 (2004). 10.1016/j.abb.2004.02.013

15 Stwora-Wojczyk, M. M., Kissinger, J. C., Spitalnik, S. L. & Wojczyk, B. S. O-glycosylation in Toxoplasma gondii: identification and analysis of a family of UDP-GalNAc:polypeptide N-acetylgalactosaminyltransferases. Int J Parasitol 34, 309–322 (2004). 10.1016/j.ijpara.2003.11.016

16 Bandini, G., Albuquerque-Wendt, A., Hegermann, J., Samuelson, J. & Routier, F. H. Protein O- and C-Glycosylation pathways in Toxoplasma gondii and Plasmodium falciparum. Parasitology 146, 1755–1766 (2019). 10.1017/S0031182019000040

17 Revoredo, L. et al. Mucin-type O-glycosylation is controlled by short- and long-range glycopeptide substrate recognition that varies among members of the polypeptide GalNAc transferase family. Glycobiology 26, 360–376 (2016). 10.1093/glycob/cwv108

18 Yakovlieva, L. & Walvoort, M. T. C. Processivity in Bacterial Glycosyltransferases. ACS Chem Biol 15, 3–16 (2020). 10.1021/acschembio.9b00619

19 Fritz, T. A., Hurley, J. H., Trinh, L., Shiloach, J. & Tabak, L. A. The beginnings of mucin biosynthesis: The crystal structure of UDP-GalNAc:polypeptide - Nacetylgalactosaminyltransferase-T1. Proc Natl Acad Sci U S A 101, 15307–15312 (2004).

20 Fritz, T. A., Raman, J. & Tabak, L. A. Dynamic association between the catalytic and lectin domains of human UDP-GalNAc:polypeptide alpha-N-acetylgalactosaminyltransferase-2. J Biol Chem 281, 8613–8619 (2006). 10.1074/jbc.M513590200

21 Kubota, T. et al. Structural basis of carbohydrate transfer activity by human UDP-GalNAc: polypeptide alpha-N-acetylgalactosaminyltransferase (pp-GalNAc-T10). J Mol Biol 359, 708–727 (2006). 10.1016/j.jmb.2006.03.061

22 Yu, C., Liang, L. & Yin, Y. Structural basis of carbohydrate transfer activity of UDP-GalNAc: Polypeptide N-acetylgalactosaminyltransferase 7. Biochem Biophys Res Commun 510, 266–271 (2019). 10.1016/j.bbrc.2019.01.084

23 Ji, S. et al. A molecular switch orchestrates enzyme specificity and secretory granule morphology. Nat Commun 9, 3508 (2018). 10.1038/s41467-018-05978-9

24 Fernandez, A. J. et al. The structure of the colorectal cancer-associated enzyme GalNAc-T12 reveals how nonconserved residues dictate its function. Proc Natl Acad Sci U S A 116, 20404–20410 (2019). 10.1073/pnas.1902211116

25 de Las Rivas, M., et al. The interdomain flexible linker of the polypeptide GalNAc transferases dictates their long-range glycosylation preferences. Nat Commun 8, 1959 (2017). 10.1038/s41467-017-02006-0

26 de Las Rivas, M., et al. Molecular basis for fibroblast growth factor 23 O-glycosylation by GalNAc-T3. Nat Chem Biol 16, 351–360 (2020). 10.1038/s41589-019-0444-x

27 Raman, J. et al. The catalytic and lectin domains of UDP-GalNAc:polypeptide alpha-N-Acetylgalactosaminyltransferase function in concert to direct glycosylation site selection. J Biol Chem 283, 22942–22951 (2008). 10.1074/jbc.M803387200

28 Elhammer, A. & Kornfeld, S. Purification and Characterization ofU DP-N-Acetylgalactosamine:Polypeptide N-Acetylgalactosaminyltransferase from Bovine Colostrum and Murine LymphomaB W5147 Cells. J Biol Chem 261, 5249–5255 (1986).

29 Pace, D. A., McKnight, C. A., Liu, J., Jimenez, V. & Moreno, S. N. Calcium entry in Toxoplasma gondii and its enhancing effect of invasion-linked traits. J Biol Chem 289, 19637–19647 (2014). 10.1074/jbc.M114.565390

30 Lira-Navarrete, E. et al. Substrate-guided front-face reaction revealed by combined structural snapshots and metadynamics for the polypeptide N-acetylgalactosaminyltransferase 2. Angew Chem Int Ed Engl 53, 8206–8210 (2014). 10.1002/anie.201402781

31. Otwinowski, Z. & Minor, W. Processing of X-ray Diffraction Data Collected in Oscillation Mode. Methods in Enzymology, C.W. Carter, Jr. & R. M. Sweet, Eds., Academic Press (New York). 276, 307-326 (1997).

32 Potterton, E., Briggs, P., Turkenburg, M. & Dodson, E. A graphical user interface to the CCP4 program suite. Acta Crystallogr D Biol Crystallogr 59, 1131–1137 (2003).

33 Winn, M. D. et al. Overview of the CCP4 suite and current developments. Acta Crystallogr D Biol Crystallogr 67, 235–242 (2011).

34 Jumper, J. et al. Highly accurate protein structure prediction with AlphaFold. Nature 596, 583–589 (2021). 10.1038/s41586-021-03819-2

35 Emsley, P., Lohkamp, B., Scott, W. G. & Cowtan, K. D. Features and development of Coot. Acta Crystallogr D Biol Crystallogr 66, 486–501 (2010).

36 Adams, P. D. et al. PHENIX: a comprehensive Python-based system for macromolecular structure solution. Acta Crystallogr D Biol Crystallogr 66, 213–221 (2010).

37 Laskowski, R. A., MacArthur, M. W., Moss, D. S. & Thornton, J. M. PROCHECK: a program to check the stereochemical quality of protein structures. Journal of Applied Crystallography 26, 283–291 (1993). 10.1107/s0021889892009944

38 Laskowski, R. A., Rulllman, J. A. C., MacArthur, M. W., Kaptein, R. & Thornton, J. M. AQUA and PROCHECK-NMR: Programs for checking the quality of protein structures solved by NMR. Journal of Biomolecular NMR 8, 477–486 (1996).

39 Chen, V. B. et al. MolProbity: all-atom structure validation for macromolecular crystallography. Acta Crystallogr D Biol Crystallogr 66, 12–21 (2010). 10.1107/S0907444909042073

40 Davis, I. W. et al. MolProbity: all-atom contacts and structure validation for proteins and nucleic acids. Nucleic Acids Res 35, W375–383 (2007). 10.1093/nar/gkm216

41 Frisch, M. J. et al. Gaussian 16, Revision A.03. Gaussian, Inc., Wallingford CT (2016).

42 Fox, B. A. et al. Type II Toxoplasma gondii KU80 Knockout Strains Enable Functional Analysis of Genes Required for Cyst Development and Latent Infection. Eukaryotic Cell 10, 1193–1206 (2011).

43 Tomita, T. et al. Toxoplasma gondii Matrix Antigen1 Is a Secreted Immunomodulatory Effector. mBio 12 (2021).

