## Supplemental Figures and Tables for "A *Toxoplasma gondii* O-glycosyltransferase that modulates bradyzoite cyst wall rigidity is structurally and functionally distinct from host homologues"

**Fig. S1**

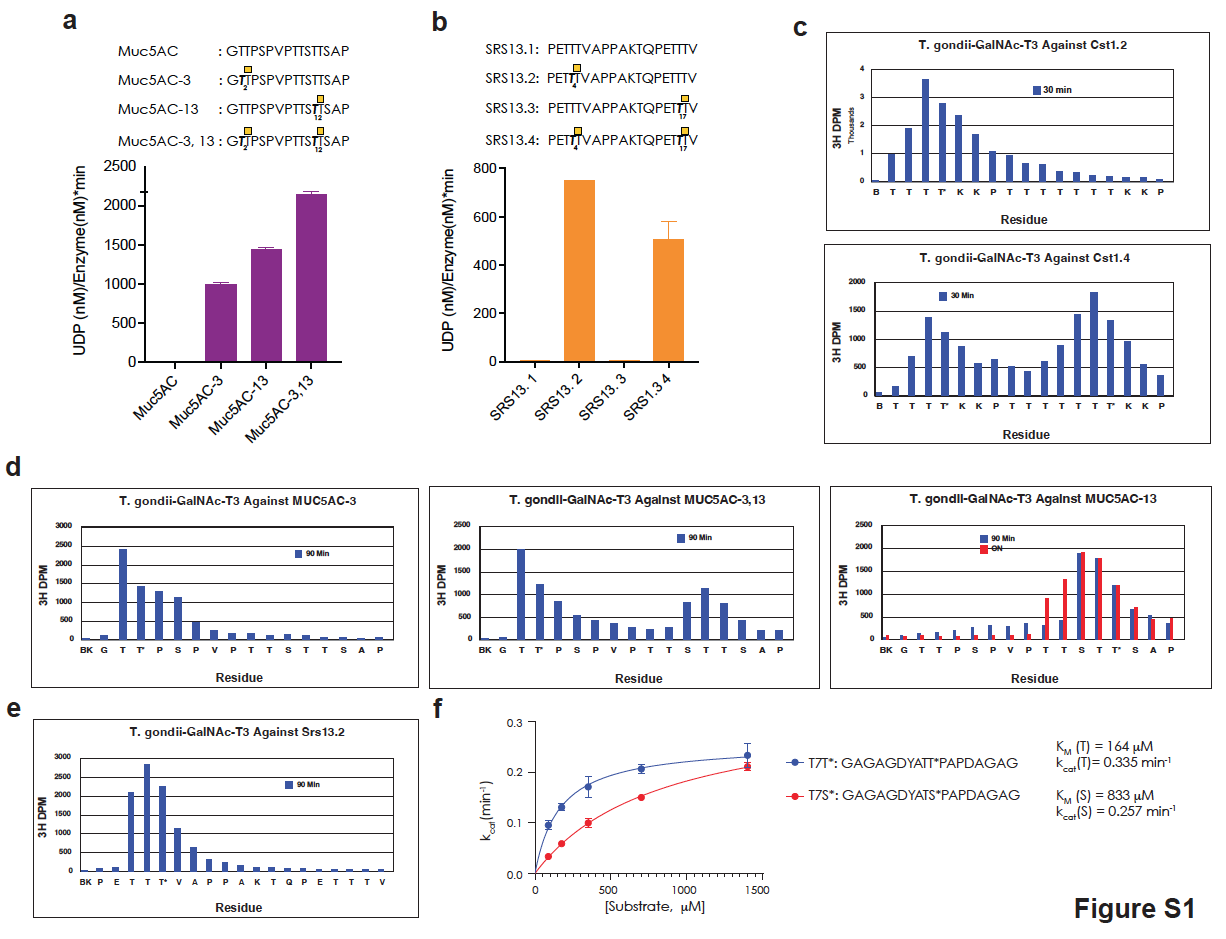

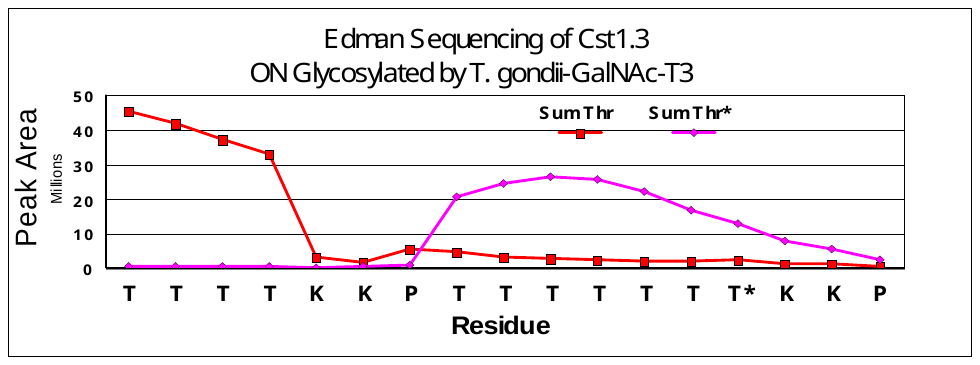

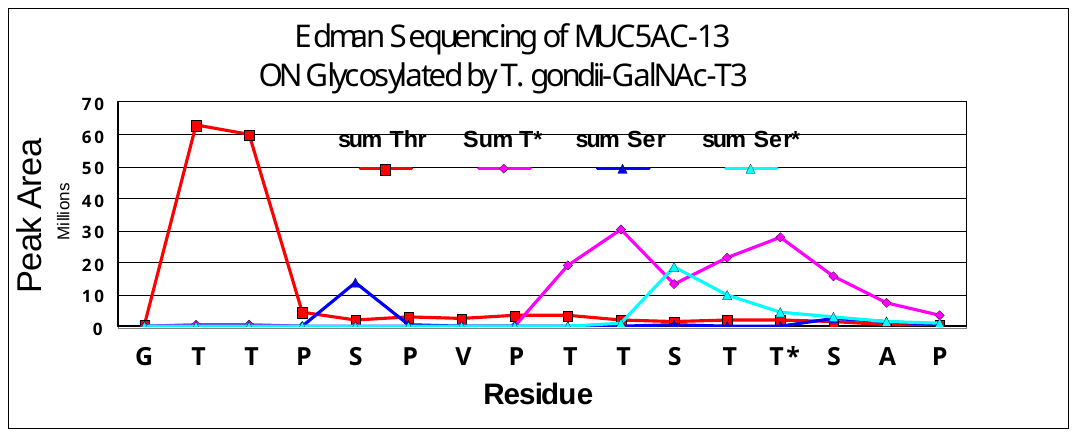
 **g**

**a,** *T.gondii*-GalNAc-T3 O-glycosylation of Muc5Ac (glyco)peptides. **b**, *T.gondii*-GalNAc-T3 O-glycosylation of Srs13 (glyco)peptides. **c,** ^3^H-GalNAc content of Edman sequencing confirming sequential and processive glycosylation of Cst1 glycopeptides after 30 min reactions and **d,** Muc5Ac glycopeptides after 90 min reaction. Like Cst1.3, the Muc5Ac-13 becomes densely O-glycosylated after an overnight reaction (right panel). **e,** Edman peptide sequencing confirm O-glycosylation of the Srs13.2 mono-glycopeptide. **f,** Activity assay comparing Thr-O-GalNAc and Ser-O-GalNAc glycopeptides shows lower K_M_ for the Thr-O-GalNAc glycopeptide, suggesting tighter binding and stronger preference for Thr-O-GalNAc over Ser-O-GalNAc for *T.gondii*-GalNAc-T3. **g**, Analysis of the Edman sequencing chromatograms of Muc5AC-13 and Cst1.3 incubated overnight with *T.gondii*-GalNAc-T3 demonstrating the nearly full glycosylation of acceptor residues N-terminal of the initial glycosylated Thr. Sum Thr represents the sum of Thr-PTH, dehydro-Thr-PTH & dehydro-Thr-DTT-PTH derivatives, sum Ser represents the sum of Ser-PTH and dehydro-Ser-DTT-PTH derivatives. Sum T* and Sum S* represents the sum of two Ser-O-GalNAc and Thr-O-GalNAc PTH diastereomers, respectively, that migrate differently on the column.

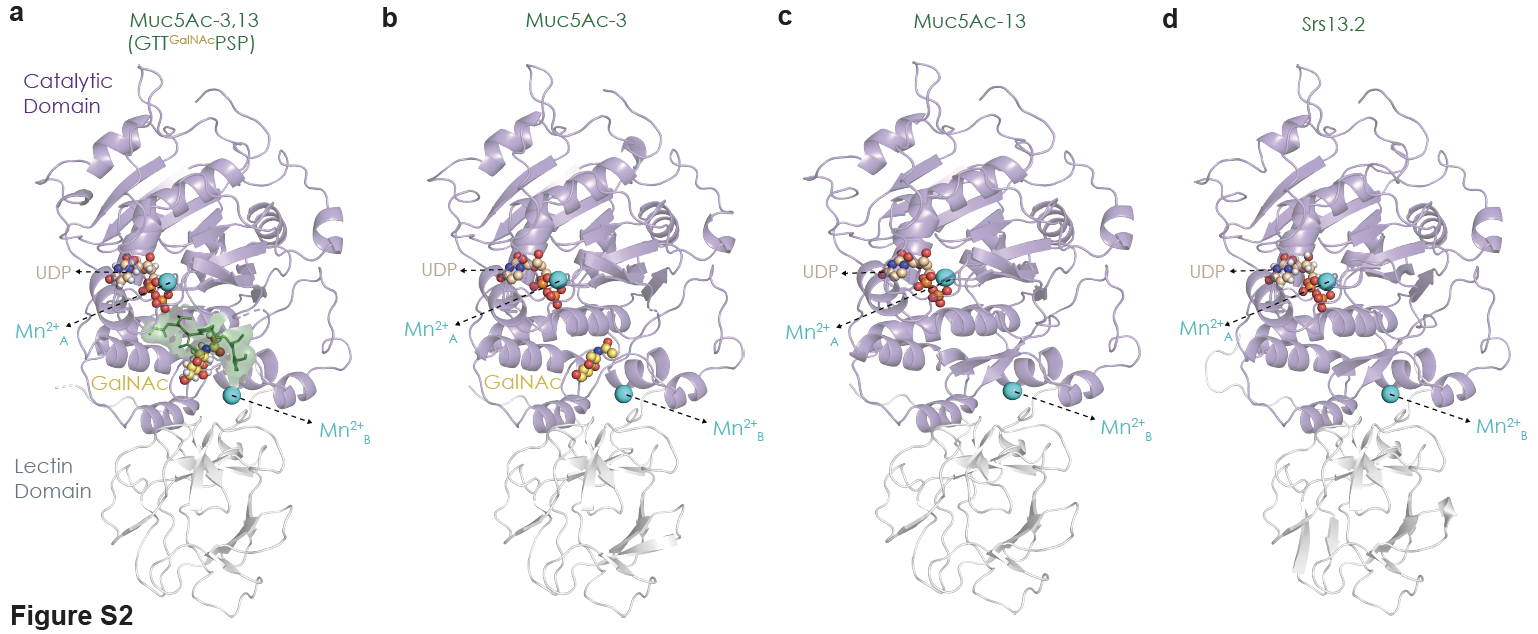
**Fig. S2**

**a-b,** X-ray crystal structures of *T.gondii*-GalNAc-T3 in complex with various mono-glycopeptides with the catalytic domain in lavender and the C-terminal lectin domain in grey. **a,** The Muc5Ac-3,13 peptide is green, GalNAc is yellow, Mn^2+^ is cyan, and UDP is wheat. Peptide density is weak in **b-d**, but GalNAc density is observed in the complex with Muc5Ac-3 suggesting the peptide is present but weakly bound in the crystal.

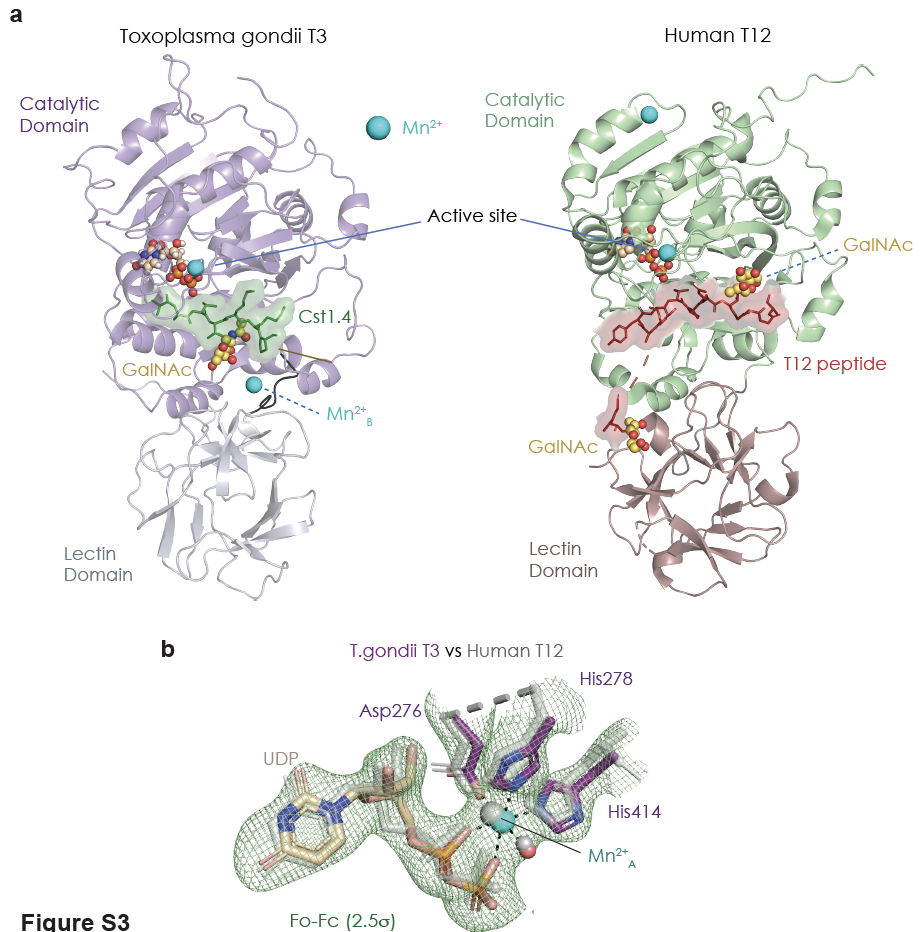
**Fig. S3**

**a,** Comparison of *T.gondii*-GalNAc-T3 co-crystal complex to the human GalNAc-T12 co-crystal complex. The overall domain architecture and active sites are conserved, with a N-terminal catalytic domain tethered to a C-terminal lectin domain by a short linker. The acceptor Thr is similarly positioned in the active site for GalNAc transfer from UDP-GalNAc. *T.gondii*-GalNAc-T3 has a second metal site (Mn^2+^_B_, shown in cyan) **b,** A superposition of the *T.gondii*-GalNAc-T3 (color) and human GalNAc-T12 (grey) active sites shows a similar configuration of the DXH motif, UDP, and Mn^2+^. The *T.gondii*-GalNAc-T3 active site Fo-Fc omit map is contoured to 2.5σ.

**Fig. S4**

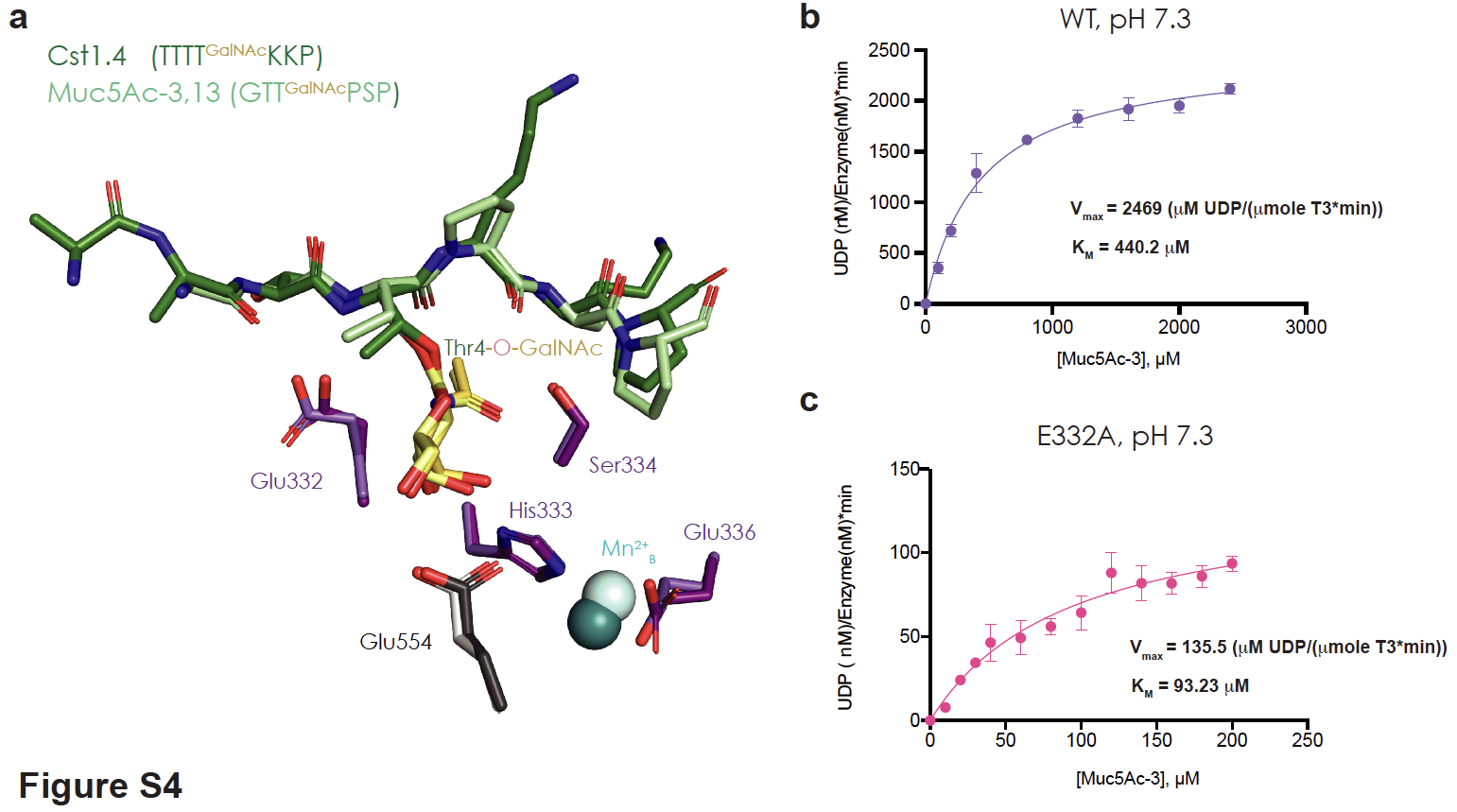

**a,** Both Cst1.4 (dark green) and Muc5Ac-3,13 (light green) have distinct peptide sequences, but similarly bind to *T.gondii*-GalNAc-T3 indicating that GalNAc binding drives substrate recognition. **b,** *T.gondii*-GalNAc-T3^WT^ kinetic plot at pH 7.3, and **c,** *T.gondii*-GalNAc-T3^E332A^ kinetic plot at pH 7.3 shows that although E332A mutation could increase binding affinity to the substrate, it primarily reduces catalytic turnover, suggesting a role in reaction chemistry.

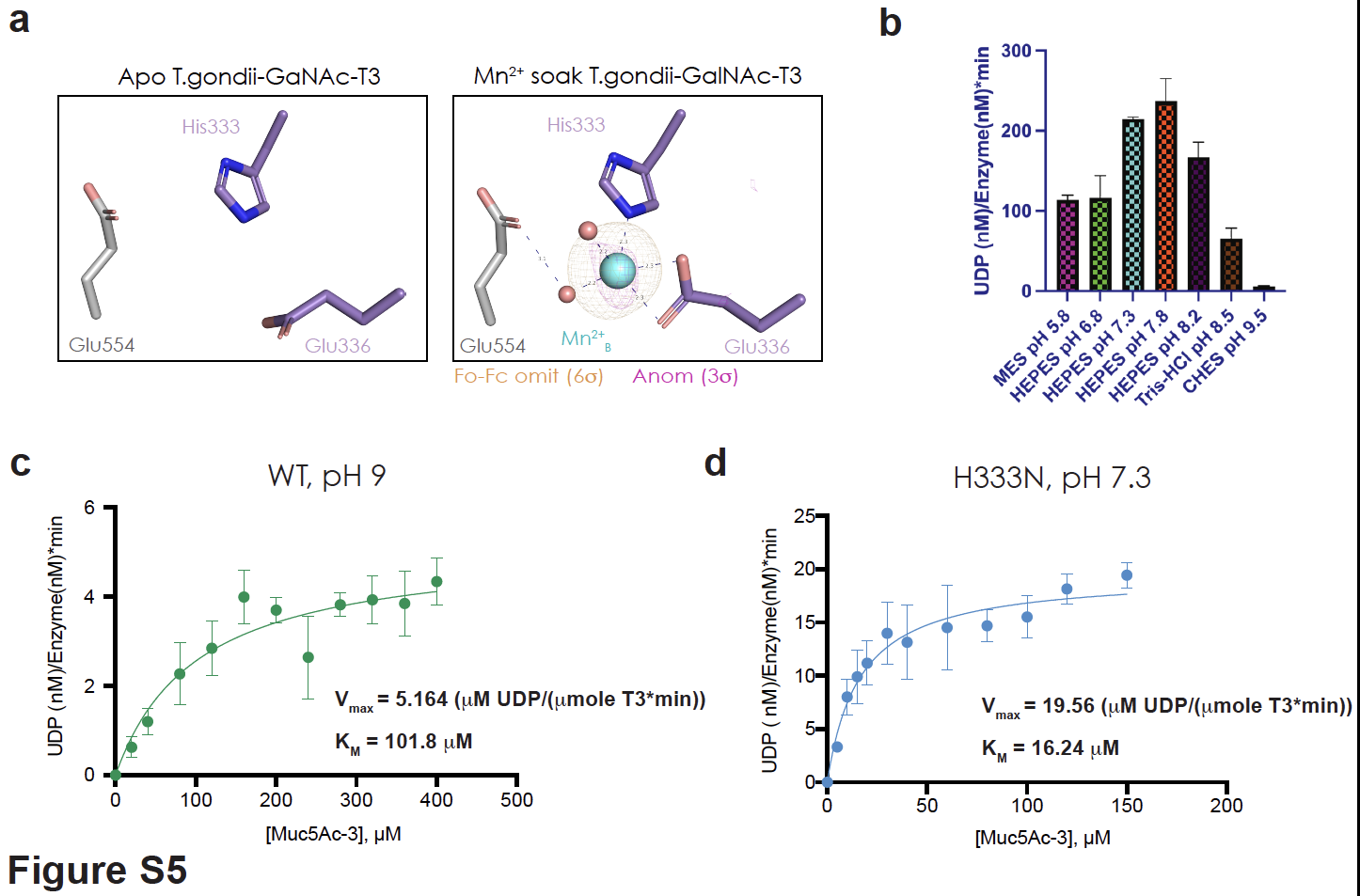
**Fig. S5**

**a,** The second metal binding residues adopt the same conformation in the absence (left panel) or presence (right panel) of Mn^2+^, where the Fo-Fc omit map is contoured to 6σ (orange) and the Anom signal is contoured to 3σ (pink). **b,** pH dependence of *T.gondii*-GalNAc-T3 activity shows optimal activity at around pH 7.3-7.8. **c,** *T.gondii*-GalNAc-T3^WT^ kinetic plot at pH 9.0 shows a ~500-fold reduction in V_max_ compared with activity at pH 7.3, with a 4-fold reduction in K_M_. **d,** *T.gondii*-GalNAc-T3^H333N^ kinetics plot at pH 7.3 shows a ~125-fold reduction in V_max_ and ~25-fold decrease in K_M_, possibly because Asn interacts more tightly to GalNAc than His.

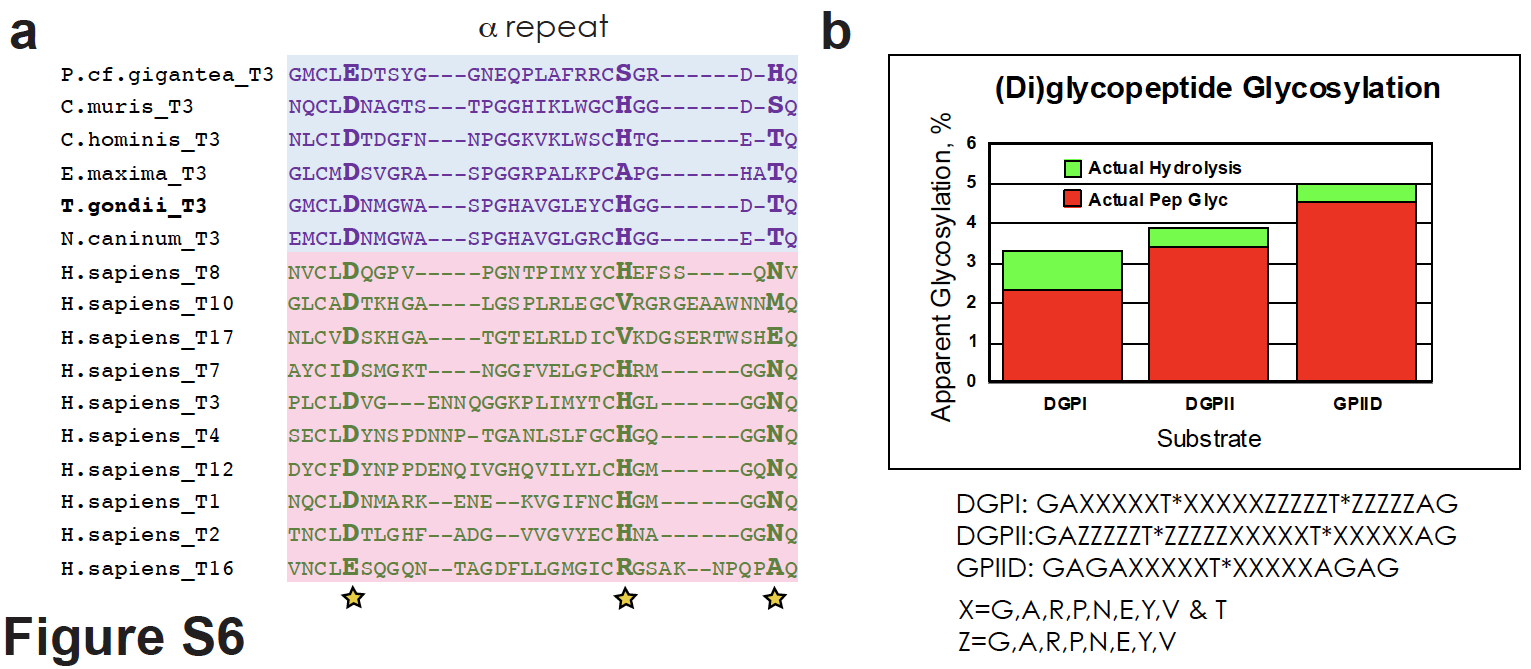
**Fig. S6**

**a,** A sequence alignment comparing the α repeat of the lectin domains of *T.gondii*-GalNAc-T3 and Apicomplexan homologues to the lectin domain of human GalNAc-Ts. *T.gondii*-GalNAc-T3 contains Asp and His required for GalNAc binding, but not Asn. Instead, this position is occupied by Thr. **b,** Probing lectin domain mediated long-range effects on activity using di-glycopeptide libraries containing GalNAc positioned N-terminal (DGPI) or C-terminal (DGPII) to the acceptor. There is no significant enhancement compared to the GPIID mono-glycopeptides, providing little evidence of lectin domain involvement in activity.

**Fig. S7**

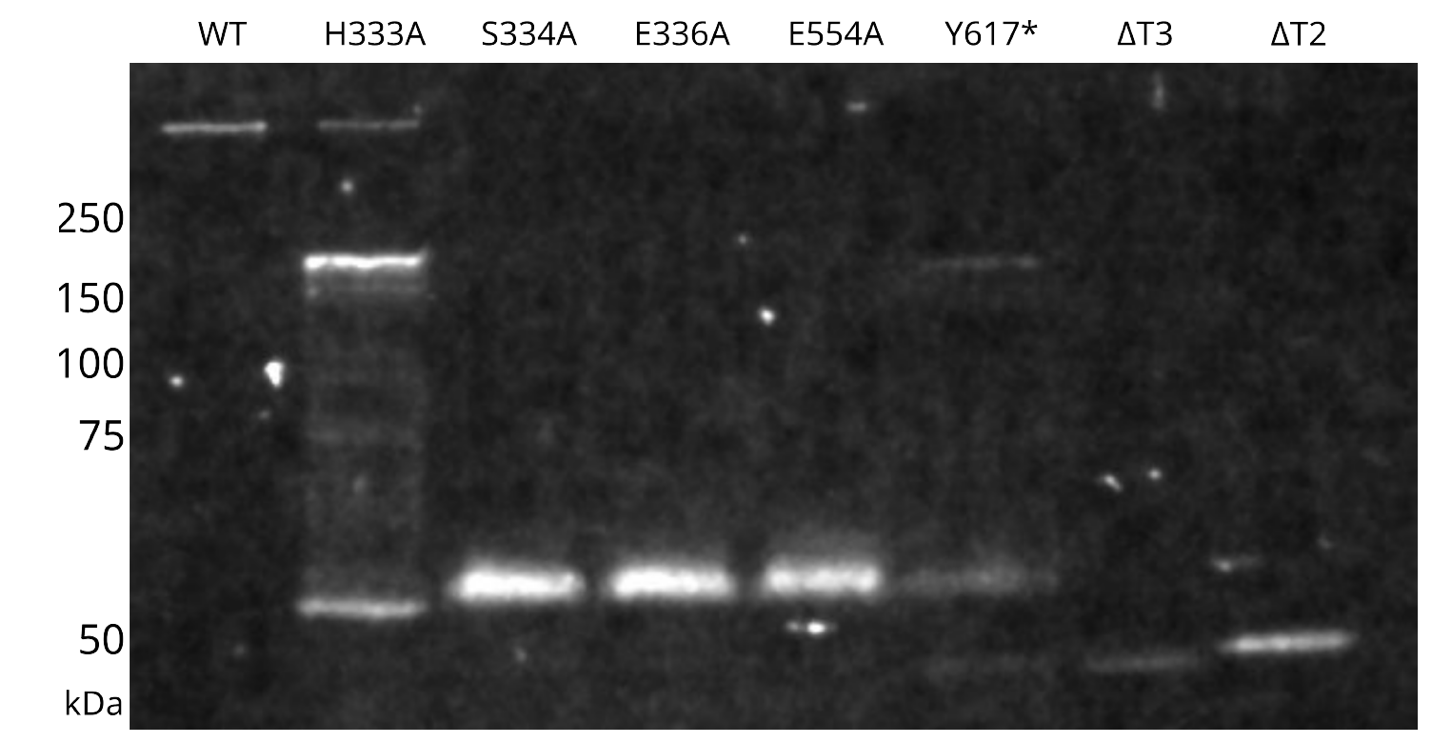

Immunoblot of the *T.gondii*-GalNAc-T3 mutant lysates probed with anti-HA antibody demonstrates the reduced glycosylation of surrogate mucin proteins in vivo. In the WT parasites, the mucin is highly glycosylated, resulting in the migration at the top of the resolving gel. In contrast, *T.gondii*-GalNAc-T3 mutants exhibited varying degrees of glycosylation deficiency in these mucin proteins.

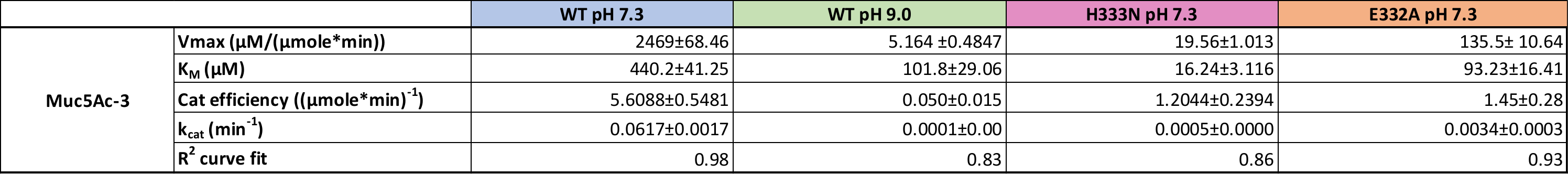
**Table S1**: *T.gondii*-GalNAc-T3 Kinetics

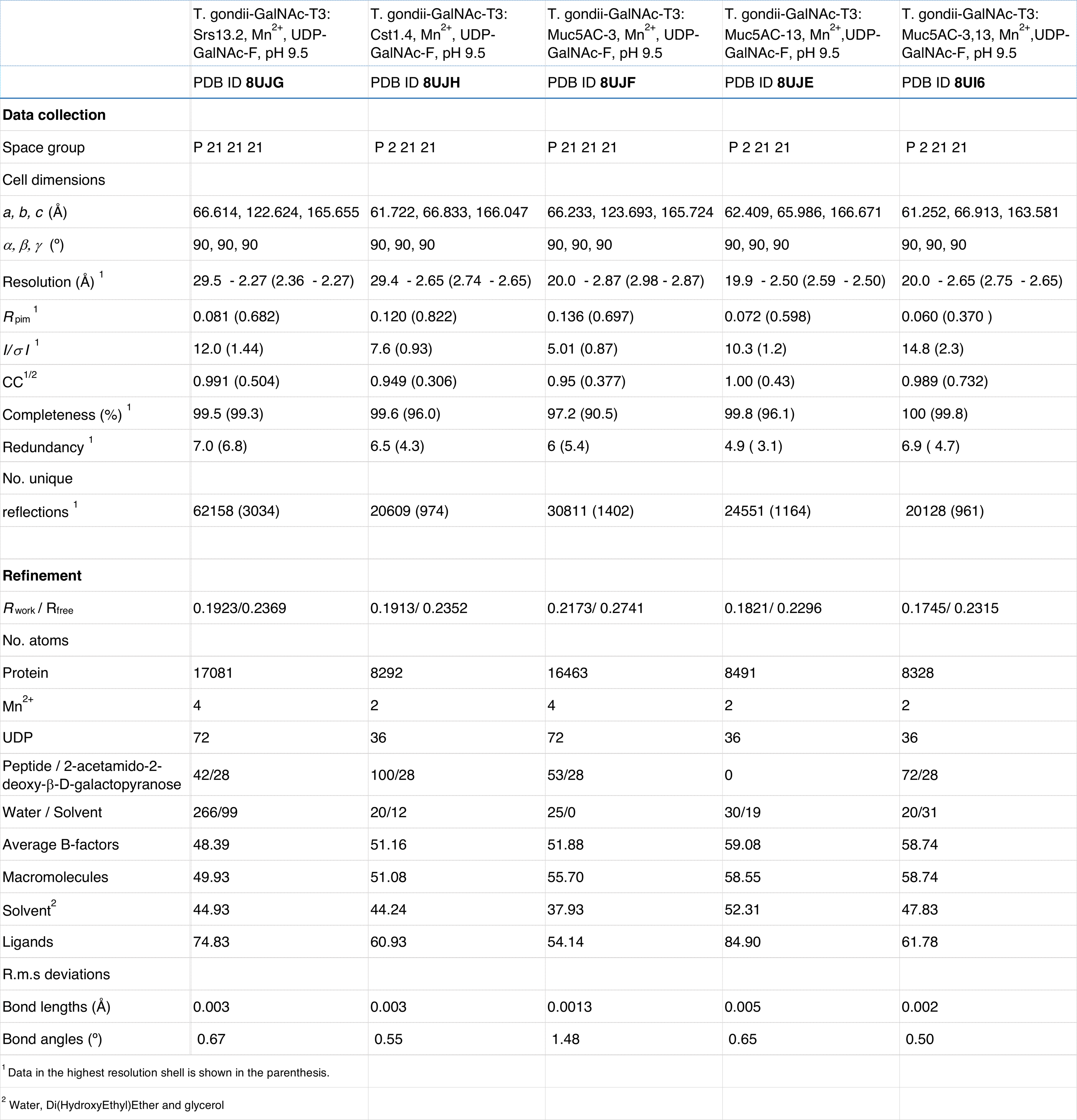
**Table S2:** X-ray and refinement statistics: Glycopeptide bound substrates

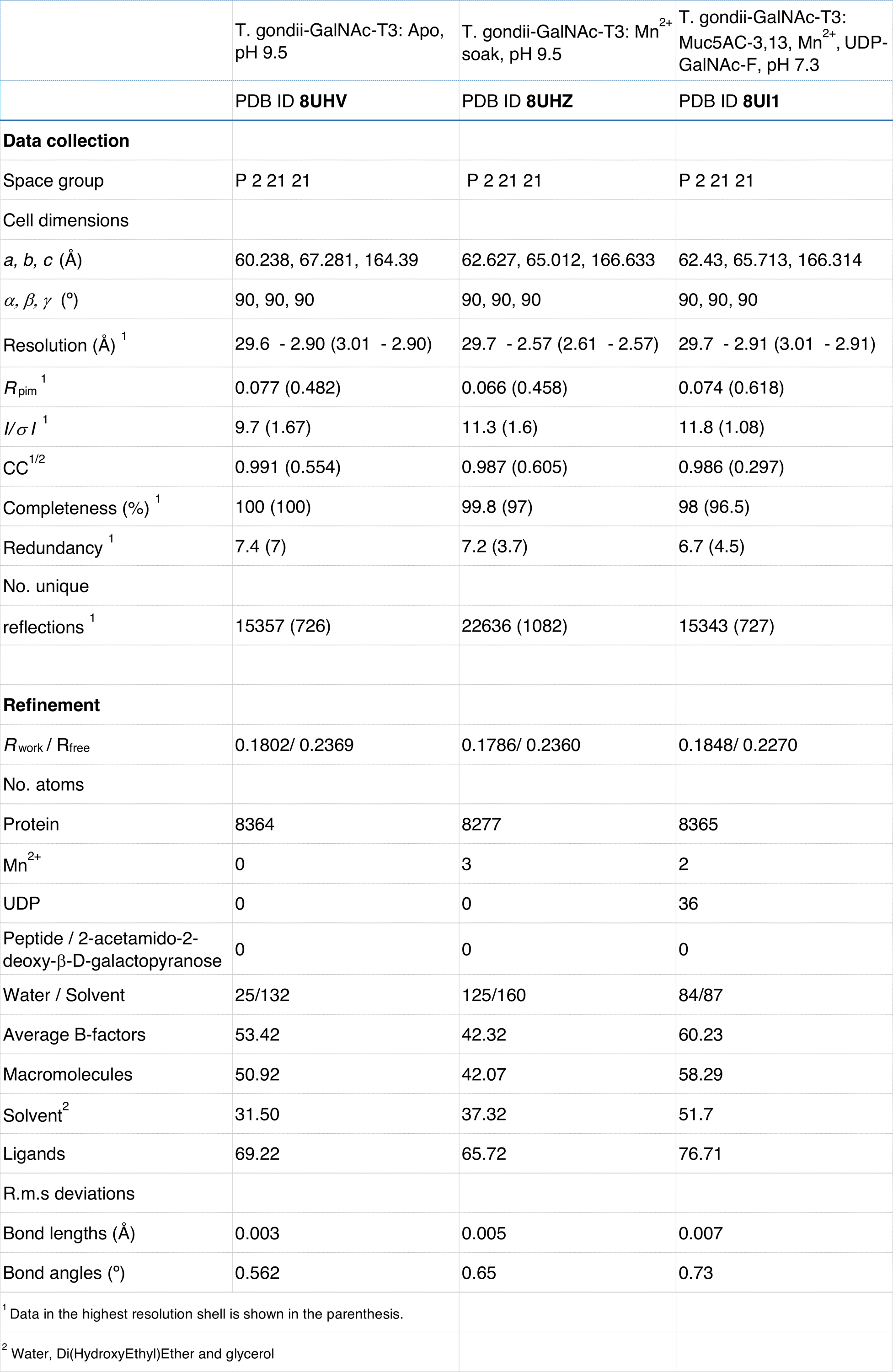
**Table S3:** X-ray and refinement statistics (Apo, Mn^2+^ soak, and pH 7.3 structures)

**Table S4:** *T.gondii*-GalNAc-T3 protein constructs and yields

| Enzyme | Approximate yield (mg/L) |
| --- | --- |
| T. gondii-GalNAc-T3 | 1.1 |
| T. gondii-GalNAc-T3_H333A | 0.020 |
| T. gondii-GalNAc-T3_H333N | 0.800 |
| T. gondii-GalNAc-T3_S334A | 0.300 |
| T. gondii-GalNAc-T3_E336A | 0.25 |
| T. gondii-GalNAc-T3_E554A | 0.800 |
| T. gondii-GalNAc-T3_ I320P | 0.300 |
| T. gondii-GalNAc-T3_ P619, P620A | 0.220 |
| T. gondii-GalNAc-T3_ F623, F625A | 0.260 |
| T. gondii-GalNAc-T3_ Y459A | 0.300 |
| T. gondii-GalNAc-T3_ E332A | 0.250 |
| T. gondii-GalNAc-T3_ E332W | 0.300 |
| T. gondii-GalNAc-T3_ E332F | 0.200 |
| T. gondii-GalNAc-T3_ E332N | 0.200 |
| T. gondii-GalNAc-T3_ E332Q | 0.200 |
| T. gondii-GalNAc-T3_ E332D | 0.150 |
